# Discovery of E6AP AZUL binding to UBQLN1/2 in cells, phase-separated droplets, and an AlphaFold-NMR integrated structure

**DOI:** 10.1101/2022.09.29.510132

**Authors:** Gwen R. Buel, Xiang Chen, Wazo Myint, Olumide Kayode, Varvara Folimonova, Anthony Cruz, Katarzyna A Skorupka, Hiroshi Matsuo, Kylie J. Walters

**Affiliations:** Protein Processing Section, Center for Structural Biology, Center for Cancer Research, National Cancer Institute, National Institutes of Health, Frederick, MD 21702, USA; Cancer Innovation Laboratory, Frederick National Laboratory for Cancer Research, Frederick, MD 21702, USA

**Keywords:** E6AP, UBE3A, UBQLN1/2, proteasome, E3 ligase, AlphaFold, liquid-liquid phase-separated droplets

## Abstract

The E3 ligase E6AP/UBE3A has a dedicated binding site in the 26S proteasome provided by the RAZUL domain of substrate receptor hRpn10/S5a/PSMD4. Guided by RAZUL sequence similarity, we test and demonstrate here that the E6AP AZUL binds transiently to the UBA of proteasomal shuttle factor UBQLN1/2. Despite a weak binding affinity, E6AP AZUL is recruited to UBQLN2 phase-separated droplets and E6AP interacts with UBQLN1/2 in cells. Steady-state and transfer NOE experiments indicate direct interaction of AZUL with the UBQLN1 UBA domain. Intermolecular contacts identified by NOESY data were combined with AlphaFold2-Multimer predictions to yield an AZUL:UBA model structure. We also identify a concentration-dependent oligomerization domain directly adjacent to UBQLN1/2 UBA (UBA-adjacent, UBAA) that is α-helical and allosterically reconfigured by AZUL binding to UBA. These data lead to a model of E6AP recruitment to UBQLN1/2 by AZUL:UBA interaction and provide fundamental information on binding requirements for interactions in droplets and cells.

## Introduction

The ubiquitin-proteasome pathway removes proteins that are misfolded or no longer needed in cells (Chen *et al*, 2020). Its substrates are marked for degradation by post-translational modification with ubiquitin (Osei-Amponsa & Walters, 2022). Ubiquitination begins with ATP-dependent charging of the ubiquitin C-terminus by an E1 activating enzyme for subsequent thioester transfer to an E2 conjugating enzyme. An E3 ligase next acts as either a scaffold to facilitate direct transfer of ubiquitin from the E2 to a substrate or as an intermediary receptor by first accepting ubiquitin from the E2 before passing it to the substrate. The E3 ligase E6AP/UBE3A belongs to the latter class of E3s and is the namesake of this protein family called HECT (homologous to the E6AP carboxyl terminus) E3s. E6AP is infamous for its roles in human disease; human papilloma viral (HPV) oncoprotein E6 binds E6AP and directs its activity towards tumor suppressor p53, contributing to cervical cancer (Huibregtse *et al*, 1993a, b; Scheffner *et al*, 1993). E6AP can also promote metastatic prostate cancer (Gamell *et al*, 2019; Paul *et al*, 2016) and is implicated in neurological disorders, with loss-of-function mutations linked to Angelman syndrome (Cooper *et al*, 2004; Kishino *et al*, 1997; Matsuura *et al*, 1997) and elevated gene dosage with autism spectrum disorders (Samaco *et al*, 2005).

An AZUL (amino-terminal z*inc*-binding *domain* of ubiquitin E3a ligase) domain (Lemak *et al*, 2011) in E6AP binds to an intrinsically disordered region in the proteasome ubiquitin receptor protein hRpn10/S5a/PSMD4, so-named RAZUL (<u>R</u>pn10 <u>AZUL</u>-binding domain) (Buel *et al*, 2020). Binding to E6AP AZUL causes RAZUL to form two α-helices that interact with two AZUL α-helices to form a 4-helix bundle and loss of this interaction leads to loss of proteasome-associated E6AP (Buel *et al*., 2020). E6AP has three isoforms with distinct localization to the nucleus or cytosol in neurons (Avagliano Trezza *et al*, 2019; Miao *et al*, 2013). Nuclear E6AP localization is contingent on AZUL-mediated interaction with hRpn10, and E6AP mislocalization causes physiological defects (Avagliano Trezza *et al*., 2019). Other E3 ligases associate with the proteasome (Besche *et al*, 2014; Buel *et al*., 2020; Leggett *et al*, 2002; Martinez-Noel *et al*, 2012; Wang *et al*, 2007) but without known binding mechanisms.

Nuclear UBE3A and proteasomes co-localize to phase-separated foci that are induced by hyperosmotic stress or nutrient deprivation and require RAD23B and ubiquitinated proteins (Uriarte *et al*, 2021; Yasuda *et al*, 2020). RAD23B and closely related Rad23A belong to a larger family of ‘shuttle factor’ proteins, so-named by their ability to deliver ubiquitinated proteins to the proteasome, that also includes DDI1/2 and UBQLN proteins (UBQLN1-4 and UBQLNL) (Walters *et al*, 2004). RAD23B and UBQLN1/2 are found associated with proteasomes purified from cells (Wang *et al*., 2007; Wang & Huang, 2008; Yu *et al*, 2016) and can stimulate proteasomal ATP hydrolysis and proteolysis (Collins & Goldberg, 2020) through a mechanism that has not yet been elucidated. UBQLN proteins can recruit an E3 ligase of unknown identity to ubiquitinate bound substrates through an interaction involving the UBA domain (Itakura *et al*, 2016), which is known to bind ubiquitin (Funakoshi *et al*, 2002; Rao & Sastry, 2002; Wilkinson *et al*, 2001) and contribute to interaction with the proteasome (Kleijnen *et al*, 2003).

The UBA in UBQLN2 contributes to UBQLN2 liquid-liquid phase separation (LLPS) (Dao *et al*, 2018; Zheng *et al*, 2021) and RAD23B’s two UBA domains similarly drive formation of phase-separated nuclear foci containing proteasomes (Yasuda *et al*., 2020). K48-linked ubiquitin chains appear to drive formation of RAD23B liquid droplets (Yasuda *et al*., 2020) and K48- or K63-linked ubiquitin chains slightly or strongly promote UBQLN2 LLPS, respectively (Dao *et al*, 2022).

Here, we find that a C-terminal region of UBQLN1/2 that includes its UBA domain has sequence similarity to the hRpn10 RAZUL, with conservation of amino acids involved in binding to E6AP. We use NMR (nuclear magnetic resonance) spectroscopy to test and confirm that the E6AP AZUL binds to the UBQLN1 UBA region. We find evidence of this interaction in cells and observe association of the E6AP AZUL with UBQLN2 phase-separated droplets in an *in vitro* assay. By integrating NMR and biophysical data with AlphaFold2-Multimer, we generate a structural model of the UBA:AZUL complex and of a UBA adjacent (UBAA) domain that is helical and self-associates. Together, our data suggest that the E6AP AZUL binds to the UBQLN1/2 UBA and this interaction allosterically impacts UBQLN UBAA self-association.

## Results

### E6AP binds to UBQLN1 and UBQLN2 in cells

Following our discovery that E6AP binds to hRpn10 RAZUL through its AZUL domain (Buel *et al*., 2020), we searched for proteins with sequence similarity to the RAZUL domain to identify other potential binders of E6AP AZUL. This approach identified a region with 34.1% and 28.6% identity (51.2% and 50% similarity) to RAZUL within UBQLN1 and UBQLN2 respectively (Figure 1A). These two isoforms are the most closely related of the UBQLN proteins (Figure S1A), with 88% sequence identity in the identified region (Figure 1A). The other UBQLN proteins were not identified in our search despite homology in this region for UBQLN3 and UBQLN4 (Figure S1B). The RAZUL helices (α1 and α2) align to a UBQLN1/2 region N-terminally adjacent to the UBA domain and to the linker between the UBA helices α1 and α2 (Figure 1A), with multiple amino acids involved in binding to AZUL conserved (Buel *et al*., 2020).

**Figure 1.**
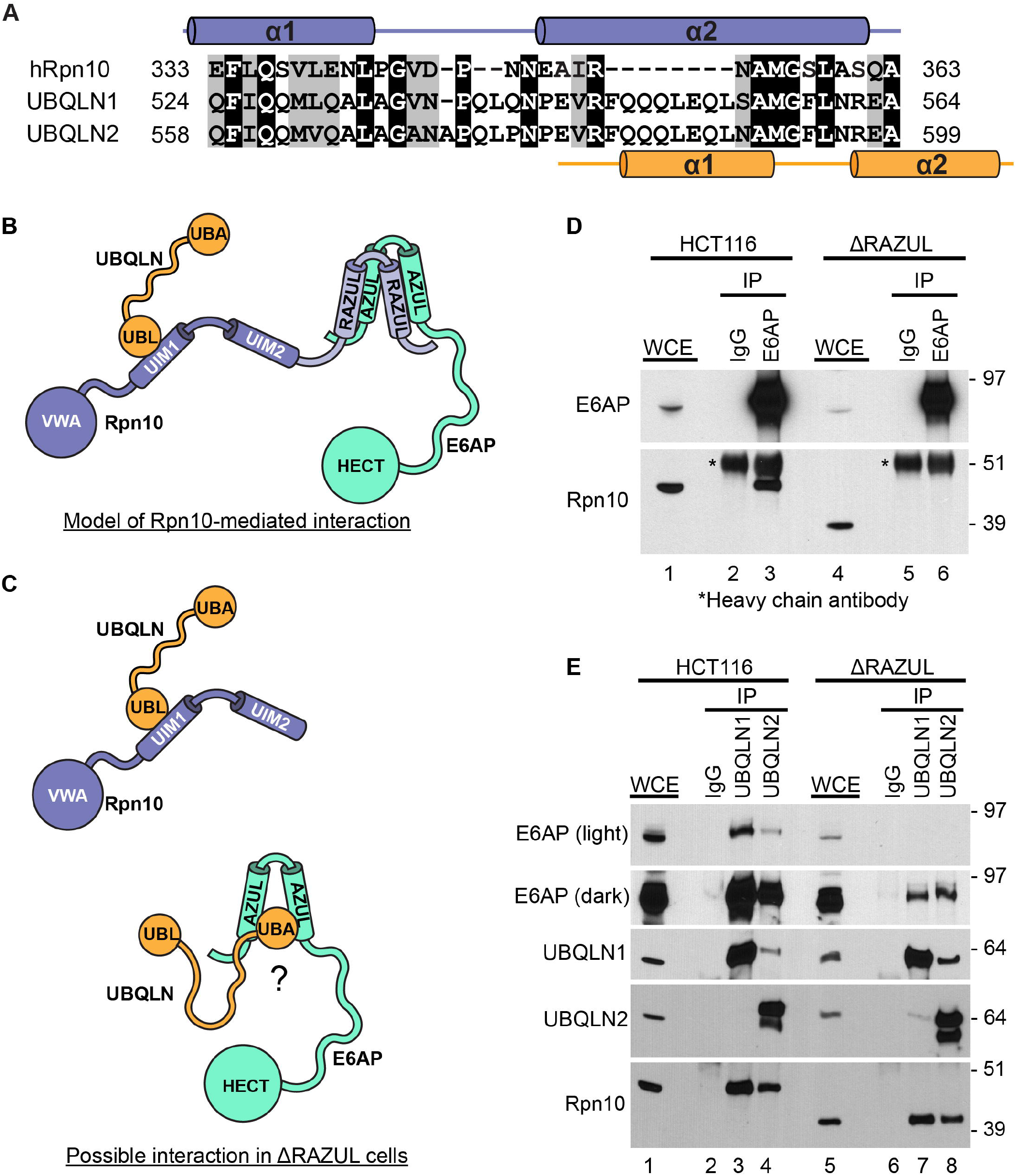
UBQLN1 and UBQLN2 interact with E6AP in cells. A) BLAST search results from uniprot.org using UniProtKB reference proteomes and Swiss-Prot databases showing sequence similarity between hRpn10 E333-A363 and a region near the C-terminus of UBQLN1 and UBQLN2. B) Model illustrating known interactions that may provide a means for E6AP and UBQLN1/2 to interact indirectly through hRpn10. C) Model illustrating how E6AP interaction with hRpn10 is lost in ΔRAZUL cells, where UBQLN1/2 interactions with hRpn10 are expected to be preserved. D) HCT116 or ΔRAZUL cells were subjected to crosslinking with DSP followed by immunoprecipitation with anti-E6AP or control antibodies. Whole cell extracts (WCE) and E6AP-immunoprecipitates were immunoblotted with the indicated antibodies. Note that hRpn10 co-precipitates with E6AP in HCT116 cells, but this interaction is lost in ΔRAZUL cells. E) HCT116 or ΔRAZUL cells were subjected to crosslinking with DSP followed by immunoprecipitation with anti-UBQLN1, anti-UBQLN2, or control antibodies. WCE and UBQLN-immunoprecipitates were immunoblotted with the indicated antibodies. E6AP is observed in both anti-UBQLN1 and anti-UBQLN2 co-precipitates in ΔRAZUL cells, albeit at lower amounts than in HCT116 cells.

To test whether E6AP interacts with UBQLN1/2 in cells, we performed immunoprecipitation experiments on cells treated with the crosslinker dithiobis(succinimidyl propionate) (DSP), an approach that allows co-immunoprecipitation of weak or transient protein complexes. One confounding issue that we anticipated for these experiments is that E6AP and the UBQLN proteins each bind to hRpn10 (Figure 1B); E6AP interacts with RAZUL (Buel *et al*., 2020) and UBQLN UBL binds hRpn10 UIMs (Walters *et al*, 2002), predominantly through UIM1 (Chen *et al*, 2019). To test whether E6AP interacts with UBQLN1/2 in an hRpn10-independent manner, we used a CRISPR-edited HCT116 cell line with RAZUL deleted (ΔRAZUL cells) (Figure 1C) (Buel *et al*., 2020). E6AP protein levels are reduced in this cell line (Figure 1D, lane 1 compared to lane 4), as demonstrated previously (Buel *et al*., 2020) by an unknown mechanism. While hRpn10 co-immunoprecipitates with E6AP in the parental cell line (RAZUL WT) (Figure 1D, lane 3), it cannot bind to E6AP when its RAZUL domain is deleted (Figure 1D, lane 6). E6AP co-immunoprecipitated with both UBQLN1 and UBQLN2 in the parental HCT116 cells (Figure 1E, lanes 3-4) as well as in the ΔRAZUL cells (Figure 1E, lanes 7-8), indicating that E6AP and UBQLN1/2 can interact independently of hRpn10 (modeled in Figure 1C). UBQLN1/2 also retained interaction with hRpn10 lacking the RAZUL domain (Figure 1E, lanes 7-8), as expected (Figure 1C).

### E6AP AZUL helices interact with UBQLN1

To test whether the E6AP AZUL directly interacts with the identified C-terminal region of UBQLN1/2, we used 2D NMR. We produced unlabeled UBQLN1 (514-589) containing a F547Y mutation (514-589, F547Y) to enable quantitation of the protein concentration by absorbance at 280 nm (Figure 1A). We added this protein to ^15^N-labeled E6AP AZUL and compared 2D ^15^N-HSQC spectra of 0.05 mM ^15^N-E6AP AZUL before and after UBQLN1 addition (Figure 2A). AZUL amide signals could be followed from free to UBQLN1-bound states (Figure 2B). To identify the specific AZUL region that interacts with UBQLN1, we quantified the signal shifting for each backbone amide signal at 1:8 molar ratio of AZUL:UBQLN1 and plotted the values according to residue number (Figure 2C) by using chemical shift assignments from our previous study (Buel *et al*., 2020). This analysis revealed the largest effects for residues in the two helices of the E6AP AZUL (Figure 2D), the same region used to bind hRpn10 RAZUL (Figure 2E).

**Figure 2.**
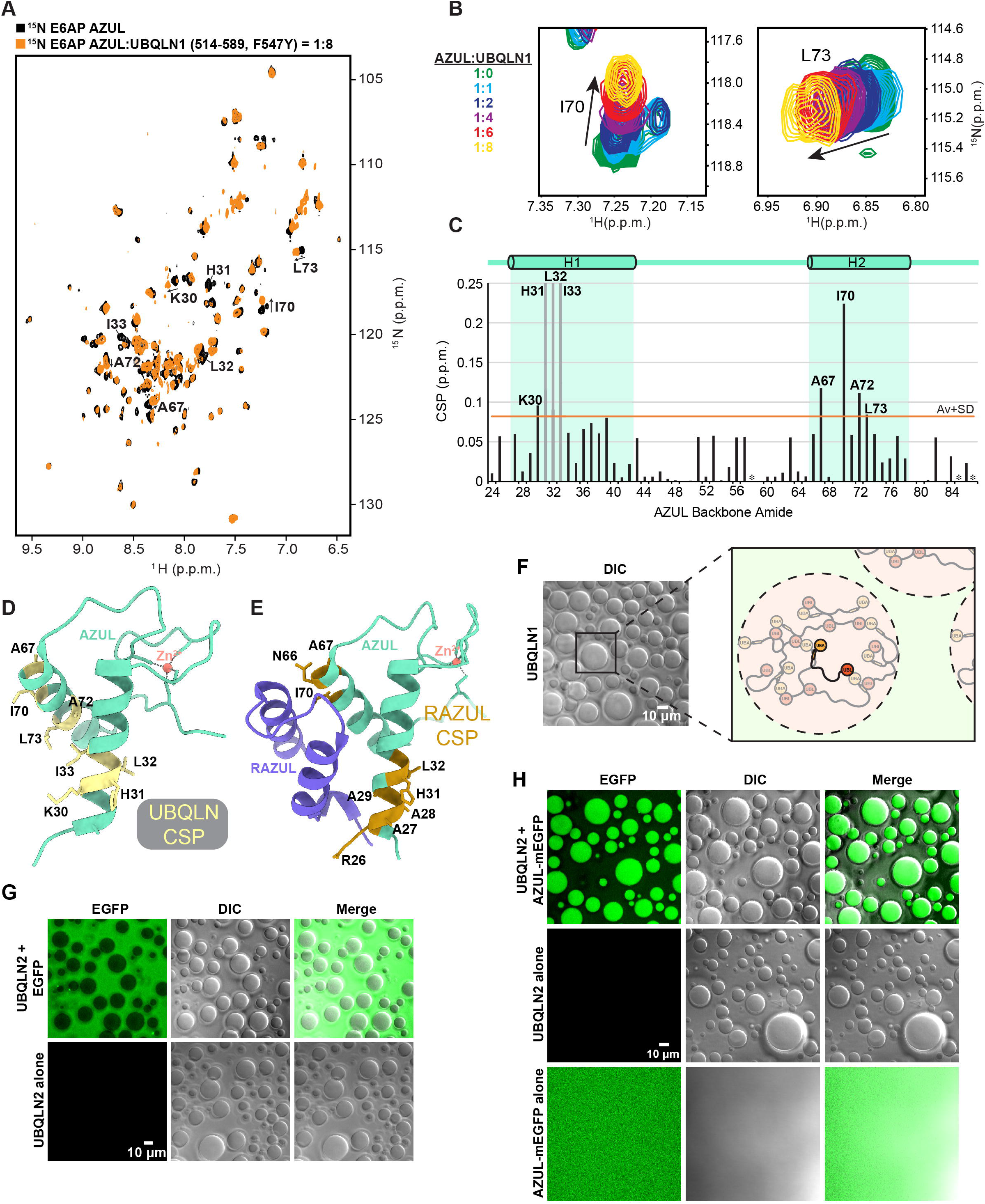
E6AP AZUL binds to the UBQLN1 C-terminal region and is recruited to UBQLN2 LLPS droplets *in vitro*. A) ^15^N-HSQC spectra of ^15^N-E6AP AZUL (black) overlayed with that of the ^15^N-E6AP AZUL mixed with 8-fold molar excess unlabeled UBQLN1 (514-589 F547Y) (orange). Peaks which either disappeared after addition of UBQLN1 or had perturbations over one standard deviation above the mean are labeled. B) Overlays of the amide signals for AZUL I70 and L73 at varying molar ratios with UBQLN1. C) Chemical shift perturbation (CSP) values derived from (A) plotted according to AZUL residue number. Gray bars indicate residues whose peaks disappeared following addition of UBQLN1. The value corresponding to one standard deviation above the mean is indicated with an orange line. D) Ribbon diagram of E6AP AZUL in which residues with CSPs over one standard deviation above the mean or having disappeared after addition of UBQLN1 are shown in yellow with side chains represented as sticks. E) Ribbon diagram of AZUL binding to the hRpn10 RAZUL (PDB 6u19) with RAZUL shown in slate blue and residues with CSPs following RAZUL addition shown in gold. F) DIC microscopy image showing 50 µM UBQLN2 in 20 mM NaPO_4_, 200 mM NaCl, and 2.5 µM ZnSO_4_ (pH 6.8) phase separating into droplets at 37 °C. Diagram to the right illustrates the high-density phase thought to exist in the UBQLN2 droplets, mediated by a large number of weak interactions between UBQLN2 monomers. G) Control experiment with UBQLN2 in the same conditions as (A) (bottom panels), or additionally containing 25 µM EGFP (top panels). Note that UBQLN2 does not fluoresce on its own (bottom left panel), and that EGFP does not colocalize with the droplets. H) 50 µM UBQLN2 and 25 µM AZUL-mEGFP in 20 mM NaPO_4_, 200 mM NaCl, and 2.5 µM ZnSO_4_ (pH 6.8) at 37 °C (top panels) or without AZUL-mEGFP (middle panels) or without UBQLN2 (bottom panels). AZUL-mEGFP alone does not phase separate (bottom panels), nor does UBQLN2 fluoresce on its own (middle panels). When mixed together, AZUL-mEGFP preferentially localizes to UBQLN2 droplets (top panels).

### E6AP AZUL is recruited to UBQLN2 droplets in vitro

UBQLN2 undergoes LLPS in cells and in isolation (Alexander *et al*, 2018; Dao *et al*., 2018), and there is some evidence that UBQLN1 may do so as well (Gerson *et al*, 2021). We therefore tested whether E6AP associates with the UBQLN proteins in the phase-separated state. We purified UBQLN1 and UBQLN2 full-length constructs and attempted to induce phase separation by incubating each sample separately at 37°C in the presence of 200 mM NaCl as done previously for UBQLN2 (Dao *et al*., 2018). Under these conditions, we observed phase separation of UBQLN2 (Figure 2F); however, no phase separation was observed for UBQLN1 (not shown). These results suggest that UBQLN1 cannot phase separate in isolation under the same conditions as UBQLN2, and perhaps different stimuli are required for UBQLN1 phase separation.

To test whether E6AP AZUL is recruited to UBQLN2 droplets, we considered various options for fluorescent tagging. Due to the presence of cysteine residues and their coordination of zinc as required for AZUL structural integrity (Kuhnle *et al*, 2018; Lemak *et al*., 2011), we avoided chemical synthesis approaches. Instead, we chose a form of EGFP to tag AZUL, since GFP-tagging had been used previously to show association of RNA polymerase II with FUS low-complexity domain phase-separated droplets (Burke *et al*, 2015) and a control experiment with EGFP alone added to UBQLN2 droplets indicated its exclusion (Figure 2G). EGFP was fused to the C-terminus of E6AP AZUL, with EGFP containing an A206K mutation (mEGFP) to decrease its propensity for dimerization (Zacharias *et al*, 2002) and avoid effects caused by unintended EGFP self-association. When we mixed AZUL-mEGFP with UBQLN2 and induced phase separation, we found the EGFP signal to localize to the UBQLN2 droplets (Figure 2H, top panels), indicating that E6AP AZUL associates with UBQLN2 in the phase-separated state. As expected, phase separation is not observed for AZUL-mEGFP in the absence of UBQLN2 (Figure 2H, bottom panels).

### The UBA-adjacent (UBAA) region of UBQLN1 is helical and structurally independent of UBA

The structure of the UBQLN UBA domain has been solved (Zhang *et al*, 2008), however there is no structural information available for UBAA. To gain insight into the structure of this region, we used AlphaFold2 (Jumper *et al*, 2021; Tunyasuvunakool *et al*, 2021) to generate a model of the UBQLN1 C-terminal region (514-589) that includes the UBAA and UBA domains (Figure 3A). The resulting structure correctly predicted the three helices of UBA (Zhang *et al*., 2008) and additionally predicted a region in UBAA to be helical (Figure 3A, residues Q524-A534). To experimentally assess the helicity of UBAA, we assigned chemical shift values to the amide H, ^H^N, C’, Cα, and Cβ atoms of UBQLN1 (514-586), as described in Methods, and inputted this information into the secondary structure prediction software TALOS+ (Shen *et al*, 2009). The UBA domain helices were identified as helical, as was the region corresponding to F525-A532 (Figure 3B), consistent with AlphaFold2 (Figure 3A). Moreover, in a ^15^N-edited nuclear Overhauser effect Spectroscopy (NOESY) experiment acquired on 0.6 mM ^15^N-labeled UBQLN1 (514-589, F547Y), we observed interactions between amide hydrogen atoms that defined the region spanning Q524-A534 as being helical (Figure S2A). Of note, no interactions were detected between UBA and UBAA, suggesting that these two regions do not interact with each other, in agreement with the AlphaFold2-predicted structure (Figure 3A).

**Figure 3.**
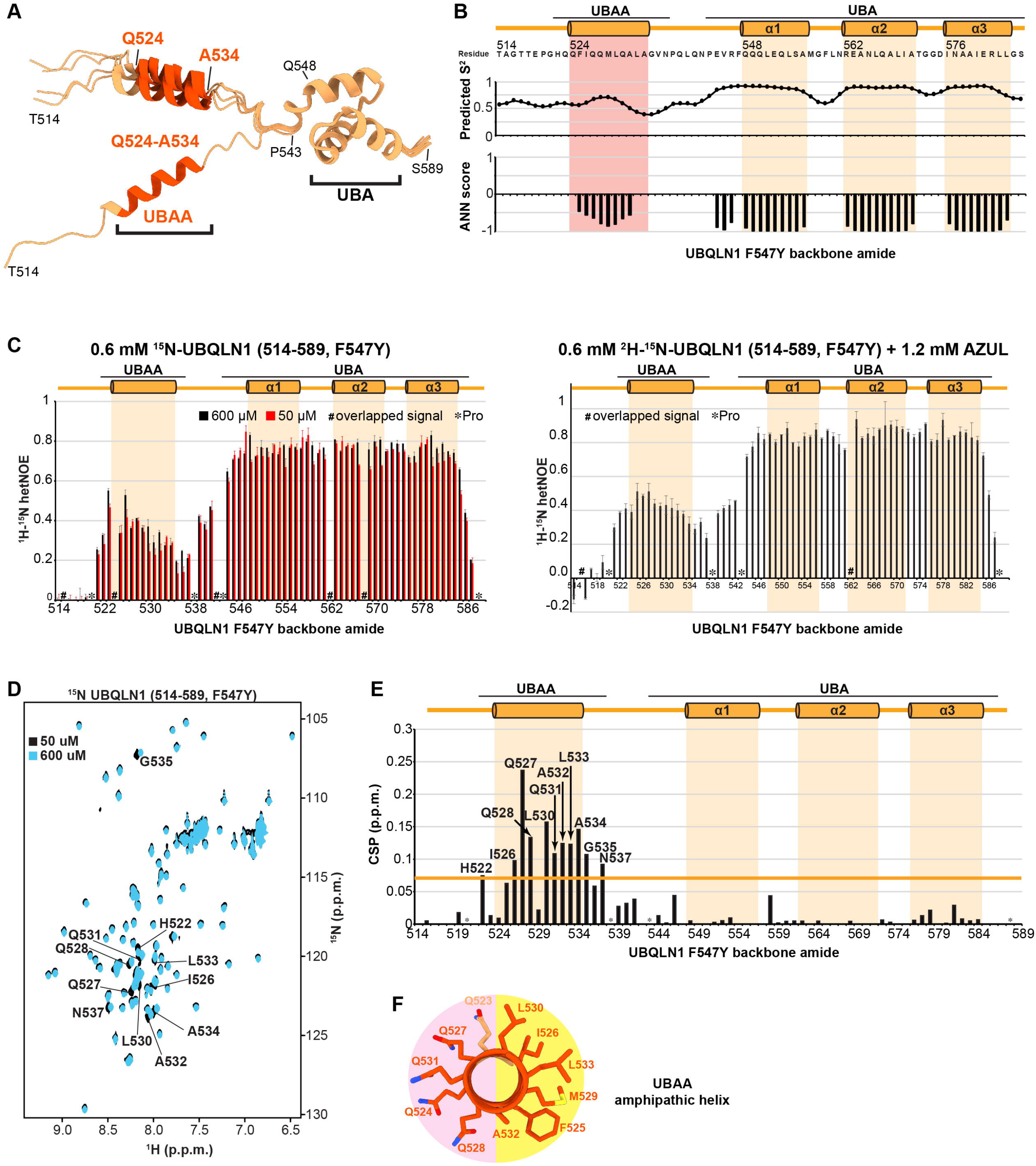
A UBA adjacent region forms a distinct, helical domain that appears to self-associate. A) AlphaFold2-predicted structures of UBQLN1 514-589 aligned based on the UBA domain. Residues predicted to be helical from data shown in (B and Figure S2A) are depicted in orange.B) TALOS+ prediction of UBQLN1 (514-586) secondary structure, with predicted order parameters S^2^ in the middle panel, and negative values in the bottom panel indicating alpha-helical prediction. Data was collected on a 0.35 mM sample. C) ^1^H-^15^N hetNOE values for UBQLN1 (514-589, F547Y) (left) and ^2^H-^15^N hetNOE values for UBQLN1 (514-589, F547Y) mixed with AZUL at a 1:2 ratio (right). D) ^15^N-HSQC spectra of ^15^N-UBQLN1 (514-589, F547Y) at 50 µM (black) and 600 µM (blue). Peaks that shift over one standard deviation above the mean are labeled. E) CSPs derived from (A) are plotted according to UBQLN1 (F547Y) residue number. The value corresponding to one standard deviation above the mean is indicated with a yellow line. F) Lengthwise view of the UBAA illustrating the amphipathy of the helix, with coloring as in (A).

As an independent assessment of whether the UBQLN1 UBA and UBAA domains interact, we measured the dynamic behavior of ^15^N-UBQLN1 (514-589, F547Y) by recording heteronuclear NOE enhancements (hetNOE). The hetNOE values in UBAA were lower than those in the UBA region (Figure 3C, left panel), with measurements at two different concentrations in agreement (600 and 50 µM). This finding indicates that these two UBQLN1 regions have distinct dynamical features consistent with UBAA and UBA not interacting, and moreover, that UBAA undergoes faster internal motion than the UBA domain.

Based on our initial hypothesis that AZUL binds to UBAA together with part of UBA (Figure 1A), we next tested whether AZUL binding could bring these two regions together. To interrogate this model, we purified ^2^H-^15^N-UBQLN1 (514-589, F547Y) and mixed it with unlabeled AZUL at a 1:2 molar ratio. The hetNOE values collected on this sample (Figure 3C, right panel) displayed similar values to those in UBQLN1 without AZUL, indicating that these domains remain separate in the presence of AZUL. In our studies of E6AP and hRpn10, we found E6AP AZUL to induce helicity in hRpn10 RAZUL (Buel *et al*., 2020). To test whether AZUL might also induce helicity in the UBQLN1 UBAA, we evaluated the circular dichroism (CD) trace of an equimolar mixture of UBQLN1 and AZUL to find a nearly identical profile compared to the theoretical sum of values obtained from UBQLN1 alone and AZUL alone, indicating that AZUL does not affect the helicity of UBQLN1 UBAA or UBA (Figure S2B).

During our NMR experiments with UBQLN1 (514-589, F547Y), we used varying protein concentrations and found that the UBAA signals varied slightly. These differences are apparent by comparing ^15^N-HSQC spectra recorded on ^15^N-UBQLN1 (514-589, F547Y) at 50 µM and 600 µM (Figure 3D). Plotting the quantified backbone amide atom shifting across the UBQLN1 sequence reveals a cluster of concentration-dependent shifts in the region spanning UBAA (H522-N357) (Figure 3E). Consistent with a model of self-association, examination of the UBAA helix from the AlphaFold2-predicted structure revealed it to be amphipathic, with a high density of glutamine residues comprising the polar side and a hydrophobic surface that could potentially drive oligomerization (Figure 3F).

### UBAA undergoes concentration-dependent oligomerization

To investigate further the dynamics of individual amino acid residues in UBQLN1 at the picosecond to nanosecond time scale, we performed NMR longitudinal (R_1_) and transverse (R_2_) relaxation experiments at high (600 µM) and low (50 µM) concentrations of ^15^N-UBQLN1. The R_1_ rates consistently indicated slightly lower values at 50 µM compared to 600 µM concentration (Figure 4A), a trend also observed for UBA R_2_ rates (Figure 4A). In UBAA however, the region spanning I526 – G535 displayed noticeably elevated R_2_ values (Figure 4A), suggesting a more ordered conformation and chemical exchange. This finding is supportive of self-association in this region and consistent with a previous study in which regions within UBQLN2 (450-624) thought to be responsible for self-association were found to undergo concentration-dependent chemical shift changes at high versus low concentration and/or differential R_2_ rates when dimerized (Dao *et al*., 2018).

**Figure 4.**
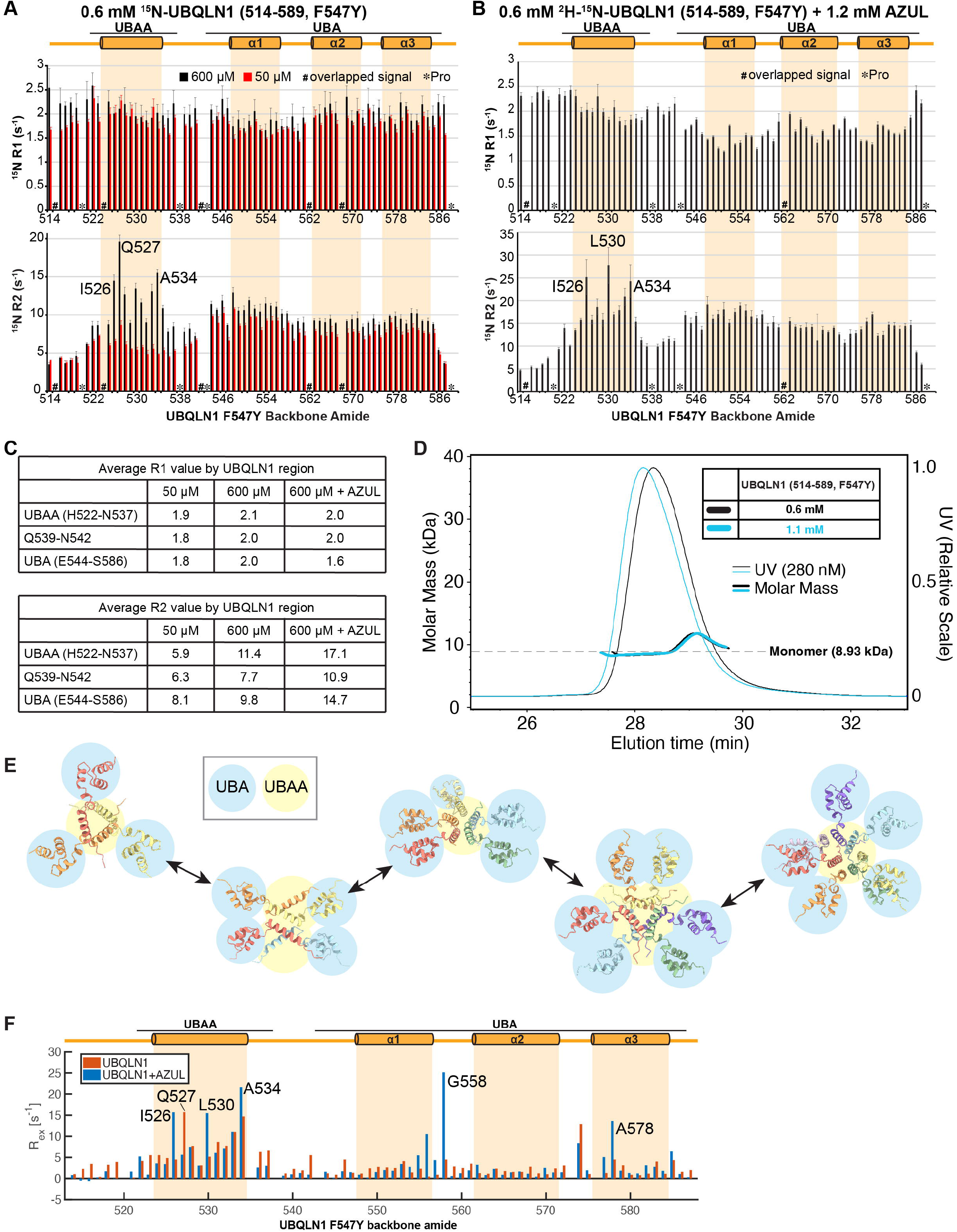
The UBQLN1 UBAA helix undergoes dynamic exchange and is more ordered at higher concentrations. A-B) ^15^N R_1_ (longitudinal) relaxation rates and ^15^N R_2_ (transverse) relaxation rates for A) ^1^H-^15^N-UBQLN1 (514-589, F547Y) or B) ^2^H-^15^N-UBQLN1 (514-589, F547Y) mixed with unlabeled AZUL at a 1:2 ratio. C) Average R_1_ and R_2_ values for UBQLN1 regions. D) SEC-MALS traces for UBQLN1 (514-589, F547Y) at 0.6 and 1.1 mM. E) AlphaFold2-Multimer predicted structures of UBQLN1 (514-589) for oligomeric states ranging from trimer to heptamer. The UBA domain of each UBQLN1 monomer is highlighted in blue, UBAA domain is highlighted in yellow. In all cases, the UBAA is predicted to cluster in the center with the UBA pointed out. F) R_ex_ values from CPMG R_2_ relaxation dispersion collected at 850 MHz with and without AZUL.

We also measured R_1_ and R_2_ relaxation rates for the same sample used in Figure 3C containing E6AP AZUL (Figure 4B). We found the R_2_ rates enhanced throughout the length of the protein compared to the sample without AZUL, as expected for deuterated samples. As in the more concentrated sample for free UBQLN1, many UBAA residues showed increased R_2_ rates compared to UBA residues. In particular, I526 and A534 were elevated in the high concentration samples, with or without AZUL present (Figure 4A-B, top panels). Q527 was elevated in the high concentration sample without AZUL, whereas L530 was elevated in the sample containing AZUL (Figure 4A versus B). R_2_ values were decreased for the region between the UBAA and UBA domains in the high concentration sample and with AZUL present, indicating it to be more flexible than the two domains under these conditions (Figure 4A-B bottom left and right panels and Figure 4C). The R_1_ rates were relatively similar throughout the length of the protein for the two concentrations of UBQLN1 (Figure 4A top panel, and Figure 4C), however, addition of AZUL reduced R_1_ values in UBA (Figure 4B top panel and Figure 4C), indicative of a larger molecular mass caused by AZUL binding.

To test further whether the concentration-dependent effects observed for UBAA are caused by self-association, we performed size exclusion chromatography coupled to multi-angle light scattering (SEC-MALS). Comparison of the UV traces between runs with different concentrations indicated the higher concentration of UBQLN1 (514-589, F547Y) to elute earlier, suggestive of self-association (Figure 4D). Furthermore, the molar mass of the eluting UBQLN1 showed two populations of UBQLN1 within the same peak (Figure 4D). The first part of the peak to elute displays a relatively flat mass profile, close to the 8.93 kDa expected mass of a monomer. However, the side of the peak that eluted later displays a slightly elevated mass (Figure 4D). The curvature of the molar masses in the later eluting side of the peak is similar to what was observed for a complex of S100B with p53 (293-393, L344P) (van Dieck, 2018) and was attributed to the dynamic equilibrium of the complex. This model is consistent with the NMR data, which suggests dynamic exchange between UBQLN1 (514-589, F547Y) monomeric and self-associated states.

We hypothesized that the UBAA portion of UBQLN1 might cluster with the hydrophobic side of its amphipathic helix occluded from the aqueous solvent. We used AlphaFold2-Multimer to generate model structures of UBQLN1 (514-589) oligomers at various stoichiometry. These structures all predict UBAA to form a helical bundle with the UBA protruding outward (Figure 4E).

To evaluate further the possibility of dynamic exchange between oligomerization states in UBAA and the effect of the AZUL domain on such exchange, we acquired CPMG R_2_ relaxation dispersion data at 850 and 600 MHz on ^2^H-^15^N-UBQLN1 (514-589, F547Y) with and without AZUL present, as chemical exchange reflects both conformational exchange as well as binding events (Figures S2A-B). As expected, field-dependent effects were observed, with greater R_ex_ (chemical exchange contribution to the apparent R_2_) at the higher field. The helical region of UBAA was found to undergo exchange in the absence of the AZUL at 850 MHz (Figure S3A) and this exchange also occurred with AZUL present (Figure S3B). To better compare the effects with and without AZUL, we plotted the 850 MHz data side-by-side for each residue to find similarity in the UBAA region, and AZUL-dependent effects in the UBA region (Figure 4F). Increased R_ex_ was observed for UBA residues when AZUL was present (Figure S3), particularly G558 and A578 (Figure 4F). These data suggest AZUL-dependent exchange in UBA and intrinsic chemical exchange in UBAA.

### E6AP AZUL interacts with UBQLN UBA

We next sought to test directly which part of UBQLN1/2 interacts with E6AP AZUL. We titrated unlabeled E6AP AZUL into ∼0.1 mM ^15^N-labeled UBQLN1 (514-586) and recorded 2D ^15^N-dispersed HSQC spectra. UBQLN1 (514-586) lacks amino acids with absorbance at 280 nm and we therefore estimated the protein concentration by 1D NMR spectra and Coomassie staining. We observed shifts in the ^1^H, ^15^N-HSQC spectrum upon addition of E6AP AZUL (Figure 5A) consistent with binding and quantified these effects for each backbone amide nitrogen and hydrogen signal in UBQLN1 (514-586). The values were plotted across the UBQLN1 sequence to identify amino acids that were significantly affected by AZUL binding (Figure 5B). We found the affected residues to map to three main areas: UBA α3, the linker between the UBA helices α1 and α2, and a residue cluster in UBAA (H527-G535) (Figure 5B). The affected residues are mapped onto a model monomeric structure in Figure 5C. We next used the two-site Carver-Richards equation (Carver & Richards, 1972) to calculate Δω values from the CPMG experiments and plotted values for residues that significantly agree with the two-site exchange model as described in Methods next to the degree of signal shifting observed in the titration experiments (Figure 5D). These analyses indicated close agreement.

**Figure 5.**
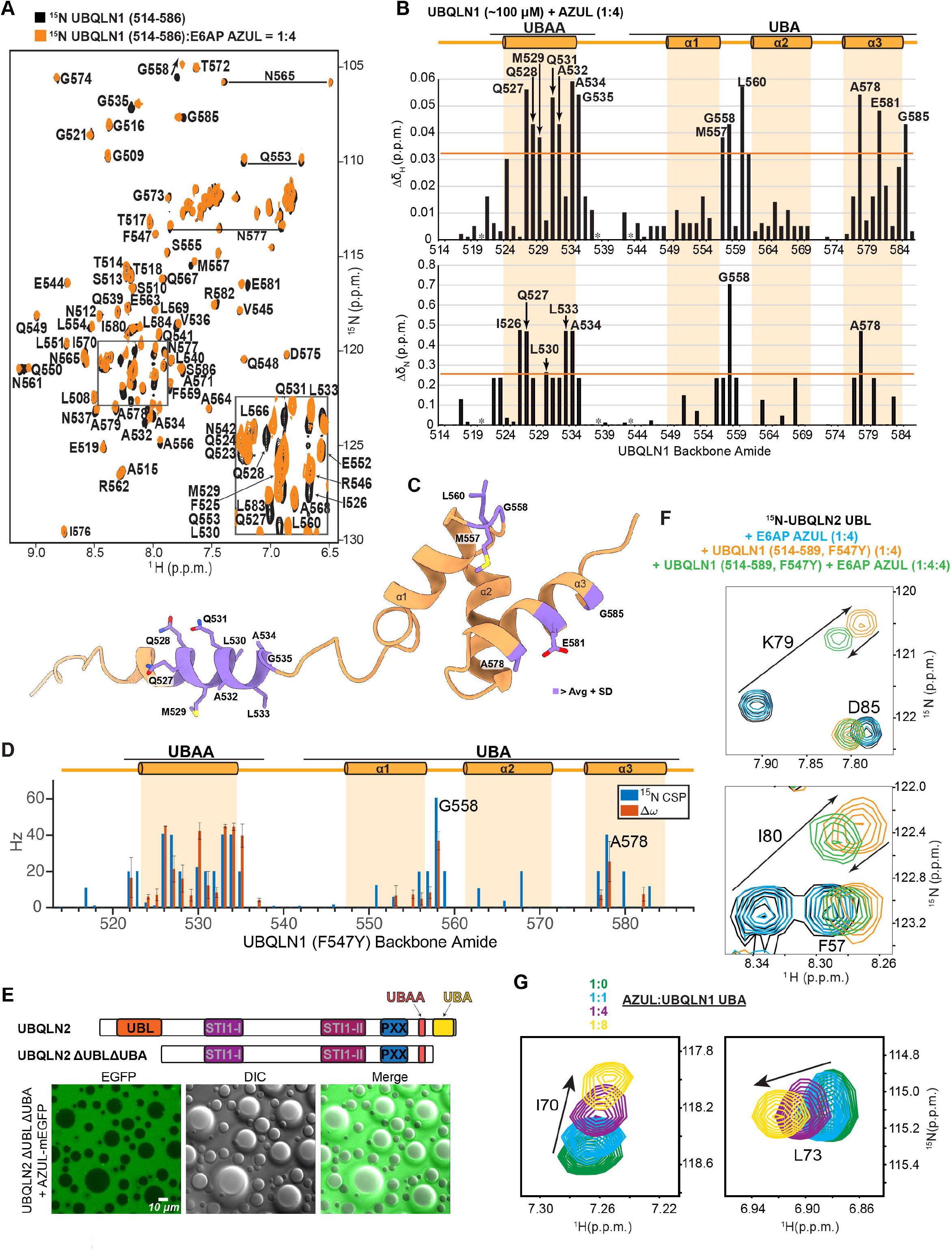
E6AP AZUL binds UBQLN UBA. A) ^15^N-HSQC spectra of ^15^N-UBQLN1 (514-586) at approximately 100 µM (black) overlayed with that of ^15^N-UBQLN1 (514-586) mixed with 400 µM unlabeled E6AP AZUL (orange). B) Changes in hydrogen (top) or nitrogen (bottom) chemical shift values derived from (A) are plotted according to UBQLN1 residue. C) Residues with chemical shift changes one standard deviation above the mean are shown in purple on the AlphaFold2-predicted structure of UBQLN1 514-589. D) CSPs from Figure 5B plotted alongside Δω values from ^2^H-^15^N-UBQLN1 (514-589, F547Y) mixed with unlabeled AZUL at a 1:2 ratio. E) Schematic of UBQLN2 domains and truncation construct (top). Lower panels show 50 µM UBQLN2 ΔUBLΔUBA and 25 µM AZUL-mEGFP in 20 mM NaPO_4_, 200 mM NaCl, and 2.5 µM ZnSO_4_ (pH 6.8) at 37 °C. AZUL-mEGFP is excluded from UBQLN2 ΔUBLΔUBA droplets. F) Zoomed views of ^15^N-HSQC spectra comparing ^15^N-UBQLN2 UBL (black) or with 4-fold molar excess unlabeled AZUL (blue), UBQLN1 (514-589, F547Y) (orange), or both AZUL and UBQLN1 (514-589, F547Y) (green). G) Overlays of AZUL I70 and L73 amide peaks from ^15^N-AZUL titrated with UBQLN1 UBA (541-589).

To directly test the importance of UBA for UBQLN interaction with AZUL, we used the UBQLN2 phase-separated droplet assay from Figure 2G-H. Initially, we wanted to remove UBA from UBQLN2 and test whether AZUL still associates with UBQLN2 droplets in its absence. However, based on previous reports, we expected UBA removal to negatively impact the ability of UBQLN2 to phase separate (Zheng *et al*., 2021). It was also reported, however, that removal of UBL in addition to UBA (i.e., ΔUBLΔUBA) increases UBQLN2 phase separation (Zheng *et al*., 2021). Therefore, we purified UBQLN2 ΔUBLΔUBA (109-576) to assess the contribution of the UBQLN2 termini in the recruitment of E6AP AZUL to UBQLN2 LLPS droplets. When AZUL-mEGFP was mixed with UBQLN2 ΔUBLΔUBA, we found the AZUL-mEGFP to be excluded from the droplets (Figure 5E), suggesting that UBA (and/or UBL) is contributing to the recruitment of AZUL-mEGFP to phase-separated UBQLN2.

We noted that the UBQLN1 UBA residues that shift upon AZUL addition were similar to those identified in UBQLN2 as interacting with UBL (Zheng *et al*., 2021). Based on these similar binding surfaces, we hypothesized that E6AP AZUL might compete with UBQLN UBL for binding to its UBA. To test this model, we purified ^15^N-UBQLN2 UBL and titrated in unlabeled E6AP AZUL, UBQLN1 (514-589, F547Y), or the two binding partners together. Addition of E6AP AZUL did not cause spectral changes to UBQLN2 UBL, indicating that the AZUL does not interact with UBL (Figure 5F). Of note, a previous report suggested that E6AP and UBQLNs might be able to interact through the UBQLN1/2 UBL (Kleijnen *et al*, 2000); this result suggests that the binding observed in the previous report is not mediated through AZUL. By contrast, UBQLN1 (514-589, F547Y) induced spectral shifting of ^15^N-UBQLN2 UBL signals, indicating binding. Addition of E6AP AZUL to the ^15^N-UBQLN2 UBL:UBQLN1 (514-589, F547Y) mixture, however, caused back-shifting of myriad UBQLN2 signals (Figure 5F), suggesting E6AP AZUL competes for the UBL binding site of the UBA domain, albeit with weaker affinity.

A previous report found the affinity of the UBQLN2 UBL:UBA interaction to be ∼200 µM (Zheng *et al*., 2021), and sequence conservation between the UBQLN1 and UBQLN2 UBA regions (Figures 1A and S1B) suggests a similar affinity for the UBQLN2 UBL:UBQLN1 UBA interaction. Since AZUL was only partially able to compete away UBA from UBL, we anticipated that the dissociation constant (K_d_) of the UBQLN1:AZUL interaction is higher than 200 µM. We used the shifting of proton amide signals (Δδ_H_) by titration of unlabeled AZUL into 50 µM ^15^N-UBQLN1 (514-589, F547Y) to estimate the K_d_ value for this interaction (Thordarson, 2011). This analysis revealed distinct values for UBA and UBAA residues; for UBA, we found the K_d_ to be ∼460 µM (Figure S4A), consistent with the competition experiment data of Figure 5F. Interestingly, even though the shifts from UBAA were larger than those from UBA in this titration, the K_d_ fitting revealed a K_d_ of ∼1 mM (Figure S4A).

As a further test that the UBA is in fact the region interacting with E6AP AZUL, we purified UBQLN1 UBA (541-589) and added it to ^15^N-AZUL. We observed chemical shift perturbations in ^15^N-AZUL upon UBQLN1 UBA addition (Figure 5G and Figure S4B) similar to those observed upon addition of UBQLN1 (514-589, F547Y) (Fig 2 A-B). Together, these data indicate that UBA interacts directly with AZUL.

### NOE interactions indicate direct binding of AZUL to UBQLN1 UBA

To unambiguously determine the AZUL:UBQLN1 binding interface, we performed a ^15^N-edited NOESY experiment on uniformly labeled ^2^H-^15^N-UBQLN1 (514-589, F547Y) mixed with 2-fold molar excess unlabeled AZUL (as used in Figures 3C, 4B, 4F and 5D). This experimental set-up detects intermolecular interactions between AZUL aliphatic and UBQLN1 amide protons (Figure S5A), as well as intramolecular interactions in UBQLN1 that involve exchangeable protons (Walters *et al*, 1997). Interactions between the UBQLN1 amide protons were readily detected as expected (data not shown), as well as those involving UBQLN1 T572 Hγ1 (Figure S5B), which likely forms a hydrogen bond to a nearby backbone carbonyl from A568, L569, or D575 (Figure S5C). Additional NOEs were detected to the backbone amides of UBQLN1 F559 and E581, as well as to the UBQLN1 sidechain amide of N577 (Figure 6A), which localize to the third helix and loop between helices 1 and 2 of UBQLN1 UBA. These regions were also implicated in AZUL binding by the CSP and exchange analyses (Figure 5A-D). We were unable to detect any NOEs to the UBAA region.

**Figure 6.**
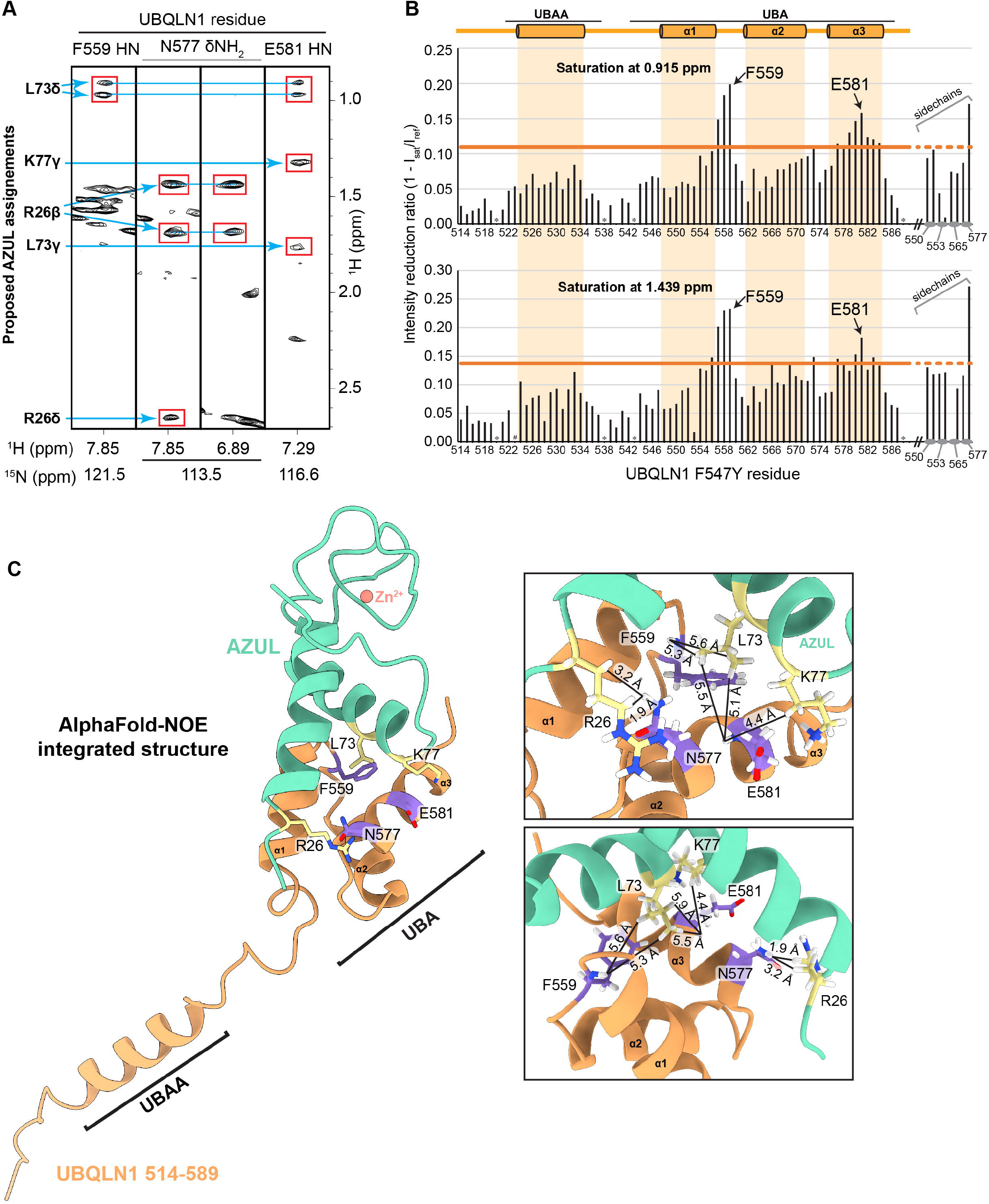
A structural model for E6AP AZUL binding to UBQLN1 UBA based on AlphaFold2-Multimer and NMR NOESY data. A) Selected regions of a ^15^N-edited NOESY collected on 0.6 mM ^2^H-^15^N UBQLN1 (514-589, F547Y) mixed with 1.2 mM unlabeled AZUL. Assignments from AZUL residues are indicated with red boxes. B) Intensity reduction ratios following saturation at 0.915 (top) or 1.439 (bottom) with the same sample as in (A). C) AlphaFold2-NOE integrated structure in which NOE assignments from AZUL were assigned guided by the AlphaFold2-Multimer-predicted structure and inputted as distance constraints into Xplor-NIH. Insets (right) show expanded regions to highlight distances between protons with assigned intermolecular NOEs. Residues from UBQLN1 with NOEs detected to their amides are shown in purple; residues from AZUL with assigned NOEs are shown in yellow; side chain nitrogen and oxygen atoms are in blue and red respectively.

The weakness and paucity of intermolecular NOEs is consistent with the weak binding affinity (Figure S4A) and to further assess binding, we performed saturation transfer experiments with the same sample. We chose saturation values based on the observed NOEs from Figure 6A (0.915 and 1.439 p.p.m.) and measured peak intensity for each UBQLN1 backbone and sidechain amide with and without the saturation. Plotting the intensity reduction ratio (1-I_sat_/I_ref_) from this analysis revealed the regions around F559 and E581 to show the greatest effect from the saturation transfer (Figure 6B), consistent with the detected steady state NOEs (Figure 6A). Most of the sidechain amides were overlapped, prohibiting their analyses; however, N577 and a few other sidechains were not overlapped. We were similarly able to observe the N577 sidechain undergoing large intensity reduction (Figure 6B). Thus, F559, N577, and E581 are at the AZUL binding interface.

### An AlphaFold-NOE integrated structure elucidates AZUL:UBQLN1 binding mechanism

We next sought to assign the intermolecular NOEs (Figure 6A) to AZUL atoms by using the deposited chemical shift assignments of apo AZUL (Lemak *et al*., 2011) and of RAZUL-bound AZUL (Buel *et al*., 2020). We were able to identify candidate AZUL atoms for the observed NOEs, however we were concerned by the likelihood that the AZUL signals might have shifted upon binding to UBQLN1. To bolster confidence in AZUL assignments, we used AlphaFold2-Multimer to predict structures of the UBQLN1:AZUL complex. Consistent with our experimental data, AlphaFold predicted binding between the AZUL and UBQLN1 UBA domain. The top scoring structures converged to a model in which the N-terminus of helix 1 in AZUL is close to N577 of UBQLN1, and helix 2 of AZUL is proximal to F559 and E581 (Figure S5D). This prediction fit our NMR data well (Figure 2A-D, 4F, 5A-D, and 6A-B), with NOEs shared by F559 and E581 mapping well to the chemical shift values of methyl groups of L73, which in the model structures is positioned between these two residues (Figure S5D). In addition, the NOEs detected to the sidechain amide of N577 matched the chemical shift values of the beta and delta hydrogens of AZUL R26, which is nearby in the model structures. We next used these intermolecular NOEs (as labeled in Figure 6A) to generate distance constraints that were used in the program Xplor-NIH to calculate structures for the AZUL:UBQLN1 complex. The structure that best matched the experimental data was similar to that predicted by AlphaFold2-Multimer, but with AZUL rotated slightly. This rotation places AZUL L73 between F559 and E581, and positions AZUL K77 closer to E581 (Figure 6C and Figure S5E).

Interestingly, in the AlphaFold-predicted UBQLN1 oligomer structures (Figure 4D), all models place UBA and the E6AP-binding site in an outward-directed orientation, with the AZUL-binding surface exposed. We modeled AZUL onto the UBQLN1 AlphaFold-predicted tetramer (Figure 7A) to find adjacent AZUL molecules experience steric clashes. We therefore inputted four UBQLN1 and AZUL molecules into AF2-Multimer to examine whether the predicted UBQLN1 conformation is altered by the presence of AZUL. The AlphaFold-predicted structures with AZUL were less convergent than the ones with UBQLN1 alone, but each was shifted in the UBAA packing to accommodate the binding of AZUL (compare Figure 7B to 7A). Based on this modeling, we expect that the observed chemical exchange in UBAA caused by AZUL addition may be driven by steric restrictions involving AZUL binding that alter the UBAA mode of self-association.

**Figure 7.**
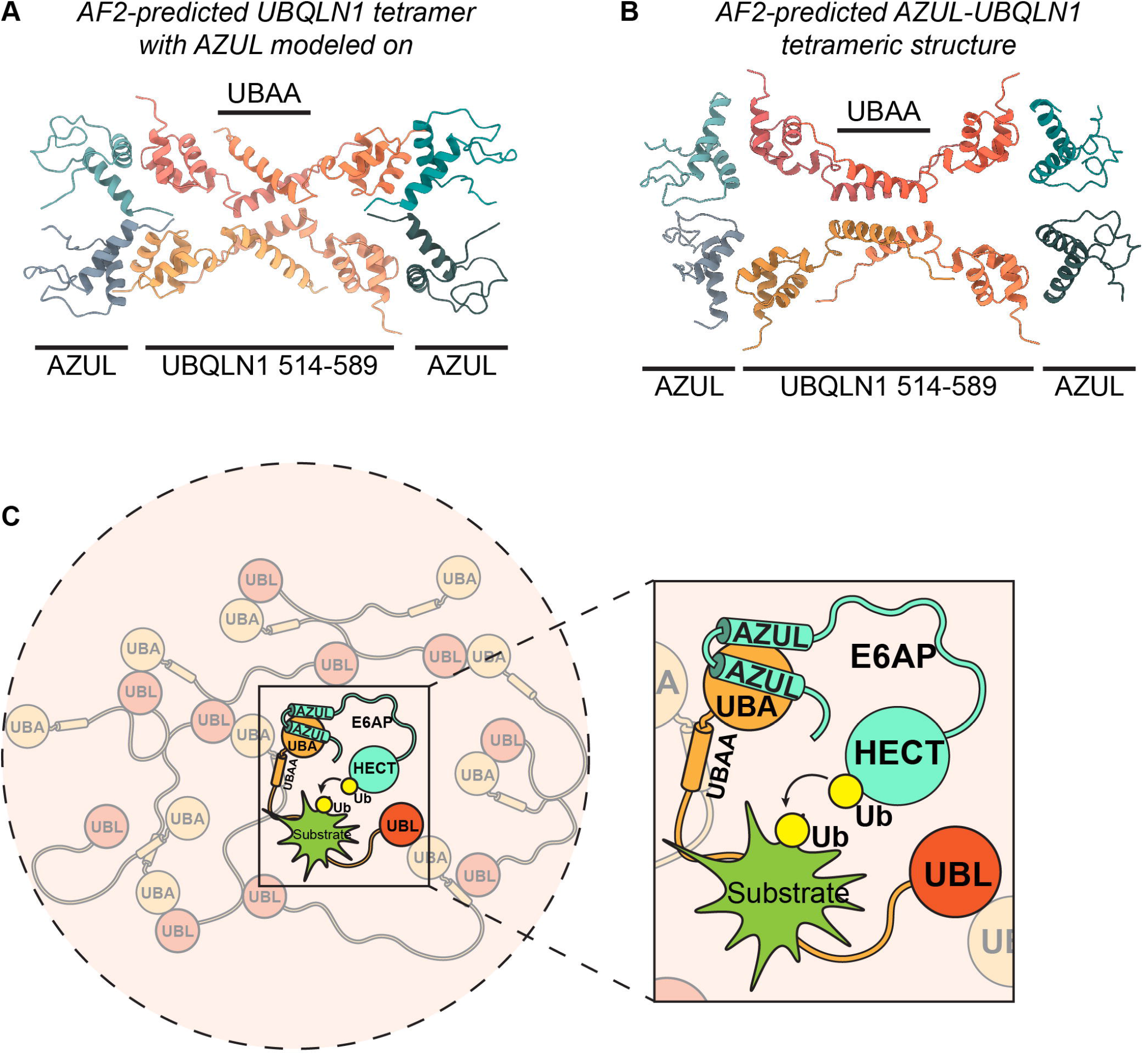
Comprehensive model of the AZUL:UBQLN1 tetramer and E6AP function in the AZUL:UBQLN1 complex. A) AlphaFold2 model of a UBQLN1 (514-589) tetramer as in Figure 4E, with AZUL modeled on based on the AlphaFold-NOE integrated structure. B) AlphaFold2-Multimer model of four UBQLN1 (514-589) and four AZUL molecules. C) Depiction of how E6AP may be interacting with UBQLN1/2 in the context of phase-separated droplets highlighting proposed E6AP ubiquitination activity on UBQLN1/2 substrates.

## Discussion

Here we show that E6AP interacts with UBQLN1 and UBQLN2 in cells and in isolation and establish a structural model of their interaction by using AlphaFold2-Multimer in combination with NMR NOESY data. We find their interaction to be weak yet UBQLN2 recruits AZUL to its droplets. Characterizing weak protein-protein interactions structurally is challenged by their dissociation while under study. Weak interactions cannot be captured by crystallization and have yet to be trapped for study by cryo-electron microscopy approaches, whereas NMR is performed in solution and highly sensitive to changes in chemical environment, including those induced by weak interactions.

We previously found that AlphaFold2 is unable to predict structural defects caused by missense mutations (Buel & Walters, 2022); however, this study shows benefits of using AlphaFold to accelerate and complement NMR data acquired on protein complexes and this application of AlphaFold may be particularly important for weakly interacting systems. We establish an NMR-AlphaFold pipeline that can be applied to other systems to accelerate NMR data analyses and overcome barriers of using NMR for structure calculations by using AlphaFold-Multimer modeling as an independent tool to aid intermolecular NOE assignments, akin to previous efforts in which AlphaFold models were used to aid the solution of crystal structures (Barbarin-Bocahu & Graille, 2022; Cramer, 2021; Ferrario *et al*, 2022; Flower & Hurley, 2021; McCoy *et al*, 2022). We outline the workflow taken to achieve our final structural model in Figure S5F.

The prospect of E6AP interacting with UBQLN2 in the phase-separated state within cells as suggested by our *in vitro* data is an intriguing one, considering that phase-separated compartments have the ability to increase local concentrations (allowing weaker affinity interactions) and enzymatic rates (Peeples & Rosen, 2021). In this model, it is possible that E6AP interacts with UBQLN proteins at known phase-separated compartments such as stress granules, however, we were unsuccessful at attempts to observe E6AP localizing to arsenite-induced stress granules (data not shown). Considering that E6AP and UBQLN2 are both enriched in brain, another possibility is that they interact predominantly in a brain-specific phase-separated compartment, such as those observed at nerve-terminals (Milovanovic *et al*, 2018; Wu *et al*, 2021). Alternatively, the fact that we were able to observe the interaction of E6AP and UBQLNs in HCT116 cells suggests that they may also interact either in the diffuse state, or in tiny, highly transient, phase-separated compartments distinct from stress granules.

Our data shows that E6AP AZUL interacts with UBQLN UBA through a mechanism that competes with UBL-UBA binding. Although the binding affinity for the AZUL:UBA interaction appears to be weaker than that of the UBL:UBA interaction, the UBL also has additional binding sites within UBQLN (Zheng *et al*., 2021) that appear to compete with its binding to the UBA. We anticipate that the highly dynamic nature of the UBQLN proteins with competing weak interactions between the various domains allows for accessibility of UBA to AZUL and that this interaction would also be enhanced by spatial co-localization. UBQLN1 binds to its substrates through a region in the middle of the protein sequence (Itakura *et al*., 2016) and we propose that E6AP AZUL binding to UBQLN UBA co-localizes this E3 for ubiquitination of UBQLN1 cargo (Figure 7C). We have not delineated in cells whether the E6AP:UBQLN interaction occurs in a low-density phase, a distinct phase-separated state, or at a concentration spectrum between these two extremes. A recent study on the effects of ubiquitin linkage type on UBQLN2 LLPS indicated that while K48-linked chains slightly promote UBQLN2 LLPS at very low Ub:UBQLN2 concentrations, most concentrations of K48-linked Ub decreased the propensity for UBQLN2 to phase separate. In contrast, K63-linked Ub had a much broader range of concentrations that promoted LLPS (Dao *et al*., 2022). As E6AP is known to catalyze K48-linked ubiquitin chains, we would expect ubiquitination of a substrate by E6AP to cause its exclusion from UBQLN2 LLPS droplets. As we expect cytosolic proteasomes to be predominantly in the diffuse state, this would liberate the Ub-substrate-UBQLN complex to bind the ubiquitin receptors of the proteasome through UBQLN UBL, effectively shuttling the substrate for degradation and perhaps further ubiquitination by RAZUL-associated E6AP (Buel *et al*., 2020) (Figure 7C).

Our data also points to steric constraints from AZUL:UBA binding leading to rearrangement of UBAA self-association/oligomerization. Of note, a recent publication found ubiquitin chain binding to the UBQLN2 UBA induces small shifts in the NMR signals of amino acids in the region of UBQLN2 homologous to UBAA, with the magnitude of shifting dependent on chain linkage type (i.e. M1, K11, K48, K63) (Dao *et al*., 2022); this finding suggests different steric hindrances from the chain type impact the amount of UBAA reorganization. Thus, either ubiquitin or E6AP AZUL binding to UBA appears to be sensed at the UBAA domain.

The role of the E6AP:UBQLN interaction in proteasome function is yet to be defined. The discovery of E6AP localized to proteasome foci in the nucleus suggests a role for E6AP in regulating phase-separated proteasome foci size or formation (Yasuda *et al*., 2020). However, it is unclear whether proteasomes and UBQLNs phase separate together or in distinct compartments. Therefore, it is also unclear whether the E6AP:UBQLN interaction has an impact on proteasomes directly or is simply a means to facilitate substrate ubiquitination. Association of an E3 ligase with the UBQLN UBA region has been previously proposed based on UBQLN1 UBA-dependent ubiquitination of a substrate in HEK cytosol or a reconstructed setting with substrate/UBQLN1 purified from rabbit reticulocyte lysate (Itakura *et al*., 2016). We propose that this E3 ligase is likely to be E6AP based on our findings reported here.

## Methods

### Cell culture, crosslinking, and immunoprecipitations

All cell lines were grown in McCoy’s 5A modified medium (Thermo Fisher Scientific 16600082), containing 10% fetal bovine serum (Atlanta Biologicals S12450), and maintained at 37 °C and 5% CO_2_. The HCT116 cell line was purchased from the ATCC (CCL-247) and ΔRAZUL (clone 13) was described previously (Buel *et al*., 2020). For crosslinking, cells were washed 2x at room temperature (RT) with PBS containing calcium and magnesium pH 7.2 (PBS/Ca/Mg) (either made by hand or Sigma D1283 diluted to 1x and pH’d to 7.2) and incubated with 2 mM DSP (ChemScene CS-0068460, first resuspended in DMSO to 25 mM, then diluted to 2 mM in PBS/Ca/Mg) for 30 min at RT. DSP was aspirated off and residual DSP was quenched through addition of 20 mM Tris (diluted from a 1 M stock) in PBS/Ca/Mg for 15 min at RT. Tris/PBS/Ca/Mg was aspirated off and cells were harvested in Nonidet P-40 lysis buffer (0.5% Nonidet P-40, 50 mM Tris pH 7.5, 150 mM NaCl, 1 mM PMSF, 5 µg/mL Pepstatin A, 10 mM Sodium Pyrophosphate, 10 mM NaF, 1 mM Sodium Vanadate, and Roche EDTA-free protease inhibitor tablet). Following 20,000 g centrifugation at 4 °C, lysate supernatants were quantitated with Pierce 660 nm protein assay reagent (Thermo Scientific 22660). For IP, 2 mg of protein was combined with 15 µL anti-UBQLN1 (Cell Signaling Technologies 14526) or anti-UBQLN2 (Cell Signaling Technologies 85509) antibodies, or 6 µL anti-E6AP (ProteinTech 10344-1-AP) antibody, or equivalent µg of control rabbit IgG (2 µg for E6AP IP or 1.815 µg for UBQLN1/2 IPs). Antibody-lysate complexes were allowed to incubate overnight at 4°C with end-over-end tumbling. 50 μL of Protein A Dynabeads (10002D) were added to the antibody-lysate mixtures and allowed to incubate for another 3 hours at 4°C with end-over-end tumbling. Bead-antibody complexes were washed 5x in Nonidet P-40 lysis buffer and eluted at 95 °C in 2x sample buffer (100 mM Tris pH 6.8, 4% SDS, 200 mM DTT, 20 % glycerol, 4 M urea, 0.0125% bromophenol blue).

### SDS-PAGE, antibodies, and immunoblots

Protein lysates and immunoprecipitates were subjected to SDS-PAGE on 4–12% NuPAGE Bis-Tris gels (Thermo Fisher Scientific NP0322) using MOPS SDS running buffer (Thermo Fisher Scientific NP0001). Proteins were transferred to 0.45 µm nitrocellulose membranes (GE/Cytiva Amersham 10600003) using NuPAGE transfer buffer (Thermo Fisher Scientific NP00061) containing 10% methanol. Membranes following transfer were blocked with 7% milk in tris buffered saline with 1% tween 20 (TBS-T). Blocked membranes were incubated overnight at 4 °C with primary antibodies diluted 1:1,000 in 7% milk in TBS-T. Membranes were washed five times in TBS-T and incubated with HRP-conjugated secondary antibodies diluted in 7% milk in TBS-T for 2 h. Following another five washes, blots were developed using WesternBright ECL-spray (Advansta K-12049-D50) and labForce HyBlot CL Autoradiography Film (Thomas Scientific 1141J52). Primary antibodies used were: E6AP (Sigma E8655), UBQLN2 (Novus Biologicals NBP2-25164), UBQLN1 (Cell Signaling Technologies 14526), and hRpn10 (Cell Signaling Technologies 3336). Secondary antibodies used were Sigma A9917 (anti-mouse, 1:5,000) and Sigma R3155-200UL (native rabbit secondary, 1:1,000).

### Protein expression and purification

The following constructs were purchased through GenScript with codon optimization for expression in *E. coli*: UBQLN1, UBQLN1 (514-586), UBQLN1 (514-589, F547Y), UBQLN1 UBA (541-589), UBQLN2, UBQLN2 ΔUBLΔUBA (109-576), and AZUL-mEGFP (E6AP isoform II 24-87 with a C-terminal mEGFP). All constructs except for UBQLN1 (514-586) and UBQLN1 (514-589, F547Y) were inserted into the pGEX-6P-1 vector between the PasI and XhoI sites, in frame with an N-terminal glutathione S-transferase (GST) and a PreScission protease cleavage site. The UBQLN1 UBA construct contains a tyrosine residue in the tag to enable quantitation at A_280_. UBQLN1 (514-586) and UBQLN1 (514-589, F547Y) were inserted into the pGEX-6P-3 vector between the EcoRI and XhoI sites. The UBQLN2 ΔUBLΔUBA (109-576) construct contains an N-terminal mCherry separated from the UBQLN2 sequence by a thrombin protease cut site; however, the mCherry was found to inhibit phase separation of the attached UBQLN2 ΔUBLΔUBA and was therefore removed via thrombin digestion after purification described below, followed by an additional run over a Superdex75 on an FPLC system to remove the mCherry. The mEGFP in frame with AZUL in the AZUL-mEGFP (separated by a TEV protease cut site) plasmid is based on the EGFP sequence, but additionally contains the A206K mutation (numbered relative to avGFP, A207K relative to EGFP). Plasmids were transformed into *E. coli* strain BL21 (DE3) (Thermo Fisher Scientific C600003) with ampicillin selection. Transformed cells were grown at 37 °C to OD_600_ of 0.5–0.6 and induced with 0.4 mM IPTG at 17 °C overnight. For AZUL-mEGFP, zinc sulfate was added at the time of induction to a final concentration of 20 μM. Bacteria were pelleted via centrifugation at 4,000 rpm at 4 °C for 40 minutes, using a Beckman Coulter J6-M1 centrifuge with a JS-4.2 rotor, and stored at -80 °C until purification. Frozen bacteria containing full-length or ΔUBLΔUBA UBQLN constructs were resuspended in Buffer A (50 mM Tris pH 8, 1 mM MgCl_2_, 1 mM PMSF, and 0.2 mg/mL DNase I and Roche Complete Mini protease inhibitor cocktail), which intentionally contains no NaCl to avoid phase separation during purification. In the case of UBQLN2 full-length protein, 0.5 mM EDTA was included in all purification buffers, but was removed by desalting with a Zeba desalting column (Thermo Scientific 89890). AZUL-mEGFP was resuspended in Buffer B (10 mM MOPS at pH 7.2, 300 mM NaCl, 5 mM 2-mercaptoethanol, 10 μM zinc sulfate, and EDTA-free protease inhibitor cocktail (Roche Diagnostics 11836170001)). UBQLN1 (514-586), UBQLN1 (514-589, F547Y) and UBQLN1 UBA (541-589) constructs were resuspended in Buffer C (20 mM at Tris pH 7.6, 300 mM NaCl, 5 mM DTT, and Roche Complete Mini protease inhibitor cocktail (Roche Diagnostics 11836153001)). Resuspended bacteria were lysed via sonication and centrifuged at 15,000 rpm (∼27,216g) for 30 min at 4 °C. Supernatants were incubated with pre-washed glutathione sepharose beads (Cytiva 17-0756-05) for 3 hours at 4 °C with a fresh protease inhibitor cocktail tablet added. Beads were washed 4-5 times in Buffer A (or B, for AZUL-mEGFP; or C, for UBQLN1 (514-586), UBQLN1 (514-589, F547Y), and UBQLN1 UBA (541-589)) and incubated overnight with PreScission protease to separate proteins from the GST tag. The on-bead PreScission protease digestion was inefficient for AZUL-mEGFP, so in this case, GST-AZUL-mEGFP was eluted with 20 mM Glutathione in Buffer D (20 mM NaPO_4_ at pH 6.8, 50 mM NaCl, 5 mM 2-mercaptoethanol, 10 μM zinc sulfate), and incubated overnight again with PreScission protease. Proteins released from GST were purified further through size exclusion chromatography on an ÄKTA pure FPLC system (Cytiva) using a HiLoad 16/600 Superdex200 or Superdex75 prep grade column in 20 mM NaPO_4_, pH 6.8 (UBQLN1, UBQLN2 and UBQLN2 ΔUBLΔUBA) or Buffer D (AZUL-mEGFP) or Buffer E (10 mM MOPS at pH 6.5, 50 mM NaCl, 5 mM DTT, and 10 μM zinc sulfate) (UBQLN1 (514-586), UBQLN1 (514-589, F547Y), and UBQLN1 UBA (541-589)). The AZUL-mEGFP required one additional incubation with glutathione beads to remove free GST.

E6AP AZUL and UBQLN2 UBL were expressed and purified as described previously (Buel *et al*., 2020; Walters *et al*., 2002). ^15^N ammonium chloride, ^13^C glucose, D-glucose-1,2,3,4,5,6,6-d_7_ (97-98%), and ^2^H_2_O (Cambridge Isotope Laboratories) were used for isotope labeling. ^2^H,^15^N-UBQLN1 (514-589, F547Y) was expressed similar to as was done previously for PUT3 (Walters *et al*., 1997), with cultures being equilibrated to 70% D_2_O for 24 hours, 100% D_2_O for 24 hours, and then 100% D_2_O with ^15^N ammonium chloride, D7 glucose and 20 mL BioExpress cell growth media (Cambridge Isotope Laboratories CGM-1000-DN) per liter added. Based on the mass from LC-MS, deuterium labeling of UBQLN1 (514-589, F547Y) was estimated to be >95%.

### NMR samples and experiments

NMR samples were prepared, namely 1) 0.35 mM ^15^N, ^13^C, 80% ^2^H-labeled UBQLN1 (514-586); 2) 0.05 mM ^15^N-labeled E6AP AZUL; 3) 0.05 mM ^15^N-labeled UBQLN1 (514-589, F547Y); 4) 0.6 mM ^15^N-labeled UBQLN1 (514-589, F547Y); 5) 0.05 mM ^15^N-labeled UBQLN2 UBL; 6) 0.1 mM ^15^N-labeled UBQLN1 (514-586); 7) 0.6 mM ^2^H-^15^N-UBQLN1 (514-589, F547Y) with 1.2 mM unlabeled AZUL.

2D ^1^H-^15^N HSQC and ^1^H-^13^C HSQC spectra and 3D HNCACB/CBCA(CO)NH, HNCO/HN(CA)CO, ^15^N-edited NOESY-HSQC (200 ms mixing time), and ^13^C edited NOESY-HSQC spectra (150 ms mixing time) were recorded on sample 1. ^15^N-edited NOESY-HSQC (120 ms mixing time) was recorded on sample 4. Samples 2, 3, 5, and 6 were used in 2D ^1^H-^15^N HSQC titrations. Sample 7 was used for ^15^N-edited NOESY-HSQC (1.8 s recycle delay and 200 ms mixing time) and saturation transfer experiments. Selective saturation of AZUL aliphatic protons in the saturation transfer experiments was performed with a 15-ms IBURP2 pulse centered at 0.915 or 1.439 ppm (2.5 s saturation duration and 3.5 s recycle delay). I_ref_ measurements were taken from a spectrum in which the 2.5 s irradiation was focused at -10 ppm. All NMR experiments were conducted in Buffer E (10 mM MOPS at pH 6.5, 50 mM NaCl, 5 mM DTT, 10 μM zinc sulfate, 1 mM pefabloc, 0.1% NaN3, and 5% ^2^H_2_O/95% ^1^H_2_O), except for 2D ^1^H-^13^C HSQC and ^13^C-edited NOESY-HSQC experiments, which were acquired on samples dissolved in ^2^H_2_O. All NMR experiments were conducted at 25**°**C. Spectra were recorded on Bruker AvanceIII 600, 700, 800, 850, or 900 MHz spectrometers equipped with cryogenically cooled probes.

All NMR data processing was performed using NMRpipe (Delaglio *et al*, 1995) and spectra were visualized and analyzed with XEASY (Bartels *et al*, 1995). Secondary structure was assessed by the TALOS+ program (Shen *et al*., 2009).

### NMR titration experiments

^1^H, ^15^N HSQC experiments were recorded on samples 2, 3, 5 and 6 (^15^N-labeled E6AP AZUL, UBQLN1 (514-589, F547Y), UBQLN2 UBL, and UBQLN1 (514-586)) with increasing molar ratio of unlabeled ligand (UBQLN1 (514-586), UBQLN1 (514-589, F547Y) or E6AP AZUL), as indicated. The amide nitrogen and hydrogen chemical shift perturbations (CSP) were mapped for each amino acid either with hydrogen and nitrogen values recorded separately, or according to Equation 1.

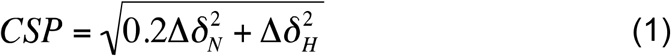

Δ*δ*_H_, change in amide proton value (in parts per million); Δ*δ*_N_, change in amide nitrogen value (in parts per million).

### NMR relaxation experiments

Rates for ^15^N longitudinal R_1_ and transverse R_2_ relaxation and magnitudes of the hetNOE were recorded on 0.05 mM and 0.6 mM ^15^N-labeled UBQLN1 (514-589, F547Y) and 0.6 mM ^2^H-^15^N-UBQLN1 (514-589, F547Y) with 1.2 mM unlabeled AZUL at 25**°**C and 700 MHz with a cryogenically cooled probe. R_1_ and R_2_ were derived by fitting data acquired with different relaxation delays (10, 20, 40, 50, 60, 80, 110, 160, 240, 320, 400, 600, 800, 1000, and 1200 ms for longitudinal relaxation; 10, 20, 30, 50, 70, 90, 110, 130, and 150 ms for transverse relaxation) to a single-exponential decay function, and error values were determined by repeating one data point. Relaxation rates were fitted by using NMRFAM-Sparky (Lee *et al*, 2015). Two spectra were recorded for steady-state NOE intensities, one with 4s of proton saturation to achieve the steady-state intensity and the other as a control spectrum with no saturation to obtain the Zeeman intensity. The control spectrum was repeated to determine error values. hetNOE were then calculated from the ratio described in Equation 2, as described in (Peng, 1994).

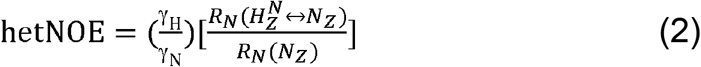

Constant Time CPMG R_2_ Relaxation Dispersion experiments were performed on 0.6 mM ^15^N-labeled UBQLN1 (514-589, F547Y) and 0.6 mM ^2^H-^15^N-UBQLN1 (514-589, F547Y) with 1.2 mM unlabeled AZUL at 25**°**C on 600 MHz and 850 MHz NMR spectrometers using the Bruker pulse sequence hsqcrexetf3gpsi3d with the total relaxation delay *Tcp* = 40 ms. Spectra were recorded with CPMG effective fields, *v*_*CPMG*_, of 25, 50, 75, 100, 150, 200, 250, 300, 400, 500, 600, 700, 800, 900, 1000, 1100, 1200, 1300, and 1400 Hz. Spectra were processed using NMRPipe and peak intensities were measured from the sum of a 3 × 3 grid centered on the peak center. Effective R_2_ rates 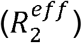 at each *v*_*CPMG*_ were determined using the relation 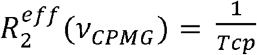 ln 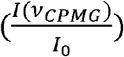 (Ishima & Torchia, 2006) where *I*(*v*_*CPMG*_) is the peak intensity after the *Tcp* relaxation period at a particular *v*_*CPMG*_ and *I*_0_ is the peak intensity without a *Tcp* relaxation period. R_2_ relaxation dispersion profiles were fit to the Carver-Richards equation (Carver & Richards, 1972; Davis *et al*, 1994; Jen, 1978) using the fmin function from the SciPy 1.7.1 package in Python 3.9. Errors in the exchange parameters were determined by using the Monte-Carlo method with a minimum of 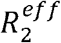error and 100 synthetic data sets.

To select only the residues from the Carver-Richards fitting that show significant chemical exchange as well as fit to a two-site model, an F-test analysis was used as previously described (Korzhnev *et al*, 2004). In brief, the R_2_ relaxation dispersion data were fit to both the Carver-Richards equation and a model that neglects exchange (optimizes for 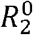at 600MHz and 850MHz only) and the F statistic was calculated for each residue. Residues were selected as undergoing chemical exchange if: 1) the F-test showed significant improvement (p < 1%) of the Carver-Richards equation over the equation that neglects exchange and 2) the 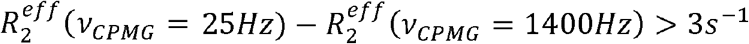. From the selected residues, the chemical shift difference between exchanging sites, Δω, was extracted and compared to chemical shift perturbation results derived from titration experiments.

### *In vitro* phase separation assays

UBQLN2 constructs and AZUL-mEGFP or EGFP were mixed at the indicated concentrations and 50 μL of the mixture was added to an Ibidi µ-Slide (Cat.No. 81506 or 81507, incubated with 3% BSA for 15 min and washed 3x with water) and imaged at 37ºC on a Zeiss LSM710 Laser scanning confocal microscope with a 63x oil, 1.4 NA, 0.19 mm objective. Images were processed using Fiji. EGFP for control experiment shown in Figure 6B was purchased from Chromotek (EGFP-250) and run through a Zeba desalting column (Thermo Scientific 89890) to exchange into 20 mM NaPO_4_ pH 6.8, 200 mM NaCl.

### Circular dichroism

Far-UV range CD spectra (260-190 nm) of 10 μM UBQLN1 (514-589, F547Y), 10 μM AZUL, and 10 μM UBQLN1 (514-589, F547Y) mixed with AZUL at a 1:1 ratio were recorded on a Jasco J-1500 CD spectrometer using a quartz cuvette with 1.0 mm path length and temperature controlled at 25 ± 0.1°C. Buffer F (10 mM MOPS at pH 6.5, 10 mM NaCl, 0.5 mM DTT, 10 μM zinc sulfate) was used as a control. All spectra were collected continuously at a scan speed of 20 nm/min and averaged over accumulation of three spectra. The buffer spectrum was subtracted from the protein spectra during data analyses. Secondary structure analysis was conducted with the program CONTIN (Provencher & Glockner, 1981; van Stokkum *et al*, 1990) by the DichroWeb server (Miles *et al*, 2022) using reference dataset SP175t (190-240 nm) (Lees *et al*, 2006).

### SEC-MALS

SEC-MALS data were collected by an Ultimate 3000 HPLC (Thermo Scientific) in-line with an Ultimate 3000 UV detector (Thermo Scientific), miniDawn MALS detector, and Optilab refractive index detector (Wyatt Technology). The data were collected following in-line fractionation with a WTC-010S5 (7.8 × 300 mm) 100 Å pore size SEC analytical column (Wyatt Technology), pre-equilibrated in Buffer E (10 mM MOPS at pH 6.5, 50 mM NaCl, 5 mM DTT, and 10 μM zinc sulfate), running at a flow rate of 0.4 mL/min. 50 μL of BSA (30 μM, Thermo Fisher Scientific 23209), or UBQLN1 (514-589, F547Y) at 0.6 or 1.1 mM was injected onto the column. Experiments were performed at 25°C. ASTRA software (version 8.0.2.5) was used for data collection and analyses.

### Electrospray ionization mass spectrometry

Mass spectrometry was performed with protein samples (∼10 μM) with 10% acetonitrile on a 6100 Series Quadrupole LC mass spectrometer (Agilent Technologies, Inc.), equipped with an electrospray source and operated in the positive ion mode. Data acquisition and analyses were performed using OpenLAB CDS ChemStation Edition C.01.05 (Agilent Technologies, Inc.).

### AlphaFold2 prediction

AlphaFold2 was utilized through the computational resources of the High-Performance Computing Biowulf cluster of the NIH (http://hpc.nih.gov). Structures were analyzed and figures generated by using PyMol (PyMOL Molecular Graphics System, http://www.pymol.org) and ChimeraX (Pettersen *et al*, 2021).

### K_d_ fitting

K_d_ estimates for the UBA and UBAA were performed using the bindfit web server (http://app.supramolecular.org/bindfit/) using the NMR 1:1 preset and the Nelder-Meid fit. Input values were Δδ_H_ (p.p.m.) values from ^15^N UBQLN1 (514-589, F547Y) titrated with unlabeled E6AP AZUL. Only residues with Δδ_H_ values greater than 0.02 p.p.m. were used for the K_d_ fitting.

## Acknowledgements

This work was supported by the Intramural Research Program through the Center of Cancer Research, National Cancer Institute, National Institutes of Health (1ZIABC011627 to K.J.W.) and in part with federal funds from the National Cancer Institute, National Institutes of Health, under contract 75N91019D00024. W.M. was supported in part by the NIH Office of Intramural Training and Education’s Intramural AIDS Research Fellowship and A.C. was supported by The Intramural Continuing Umbrella of Research Experiences (iCURE) program. This work used computational resources of the High-Performance Computing Biowulf cluster of the NIH (http://hpc.nih.gov). We thank the Optical Microscopy and Image Analysis Lab (OMAL) of NCI-Frederick for training, technical discussions, and use of microscopes. We thank Janusz Koscielniak for maintenance of the NMR spectrometers, and the Biophysics Resource in the Center for Structural Biology, CCR for assistance with LC-MS and CD-spectroscopy. We thank Dr. Angela Gronenborn for suggesting saturation transfer experiments.

## Author Contributions

G.R.B., W.M., and K.J.W. conceived of and designed the experiments. G.R.B., X.C., and O.K. performed experiments and data analyses. W.M. performed CPMG R_2_ relaxation-dispersion experiments and analyses. V.F. expressed and purified UBQLN1 UBA (541-589). A.C. assisted with sample preparation. K.A.S. assisted with SEC-MALS technical issues and data interpretation. G.R.B., H.M., and K.J.W interpreted results. G.R.B. and K.J.W. wrote the manuscript with input from all authors.

