## Supplement for "Discovery of E6AP AZUL binding to UBQLN1/2 in cells, phase-separated droplets, and an AlphaFold-NMR integrated structure"

A

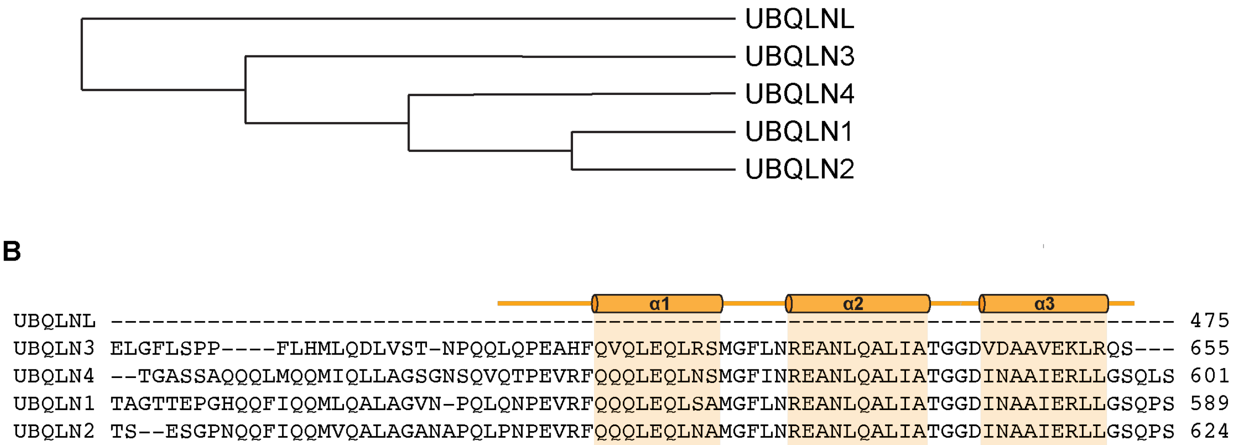


Figure S1. Related to Figure 1. A) Cladogram showing the similarity between UBQLN isoforms. B) Sequence alignment of UBQLN isoforms showing the regions homologous to UBQLN1 514-589. UBQLNL is truncated relative to the other UBQLN isoforms and does not contain this region. Alpha-helices in UBQLN UBA are shaded and indicated in orange. Both A and B were produced using Clustal Omega.

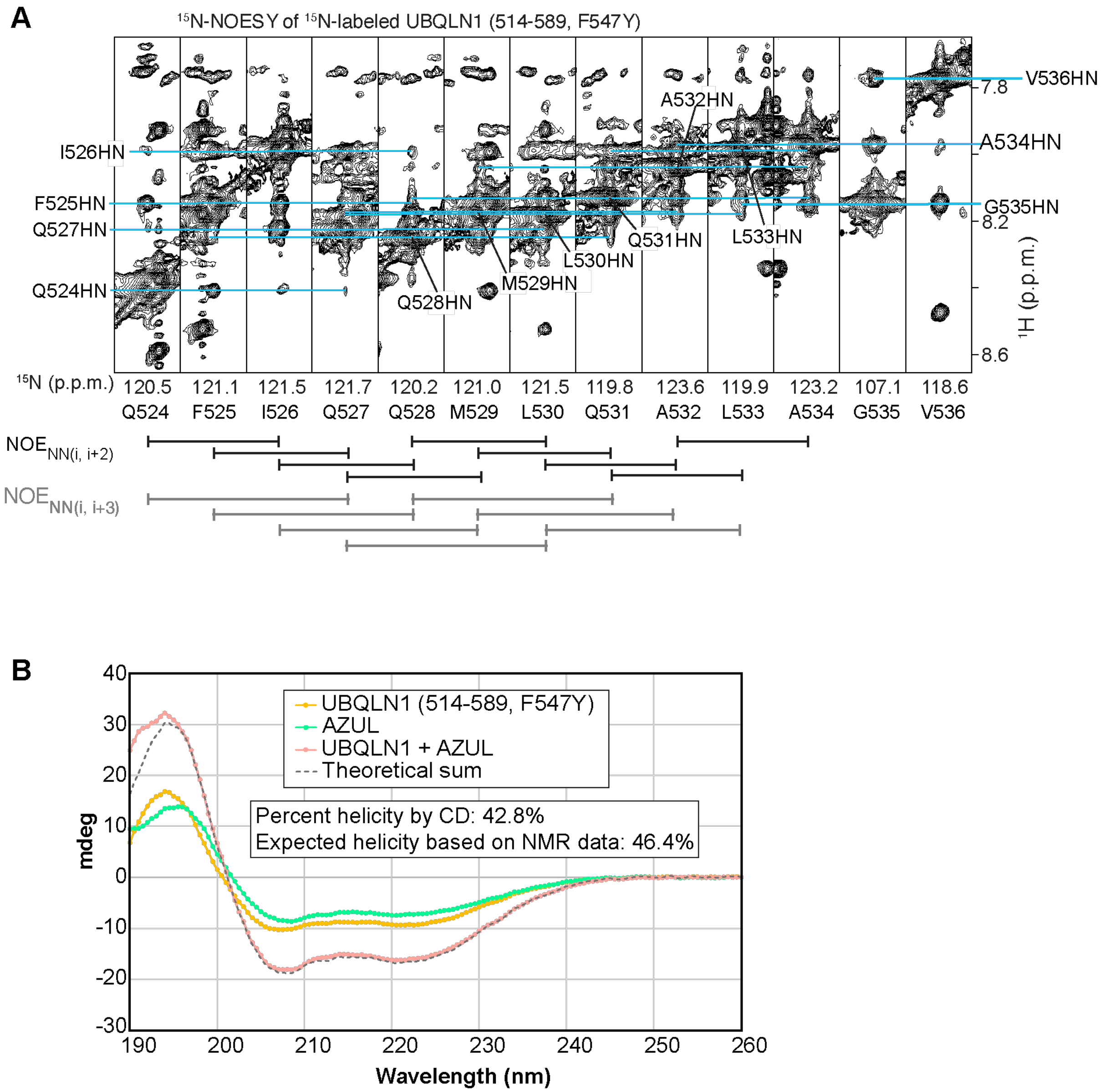


Figure S2. Related to Figure 3. A) Selected regions from a 3D $^{15}$N-dispersed NOESY spectrum acquired on 0.6 mM $^{15}$N labeled UBQLN1 (514-589, F547Y). NOEs observed between backbone amide hydrogen atoms of residues two or three residues apart (*i, i+2* and *i, i+3*) are indicated below the spectra. Blue lines are used to connect cross peaks with amino acid assignments for the indirect $^{1}$H dimension whereas the direct $^{1}$H and $^{15}$N dimension assignments are listed below the strips. B) Circular dichroism traces for UBQLN1 (514-589, F547Y), AZUL, and UBQLN1 (514-

589, F547Y) mixed with AZUL at equimolar ratio. The theoretical spectrum based on free values (gray) is displayed.

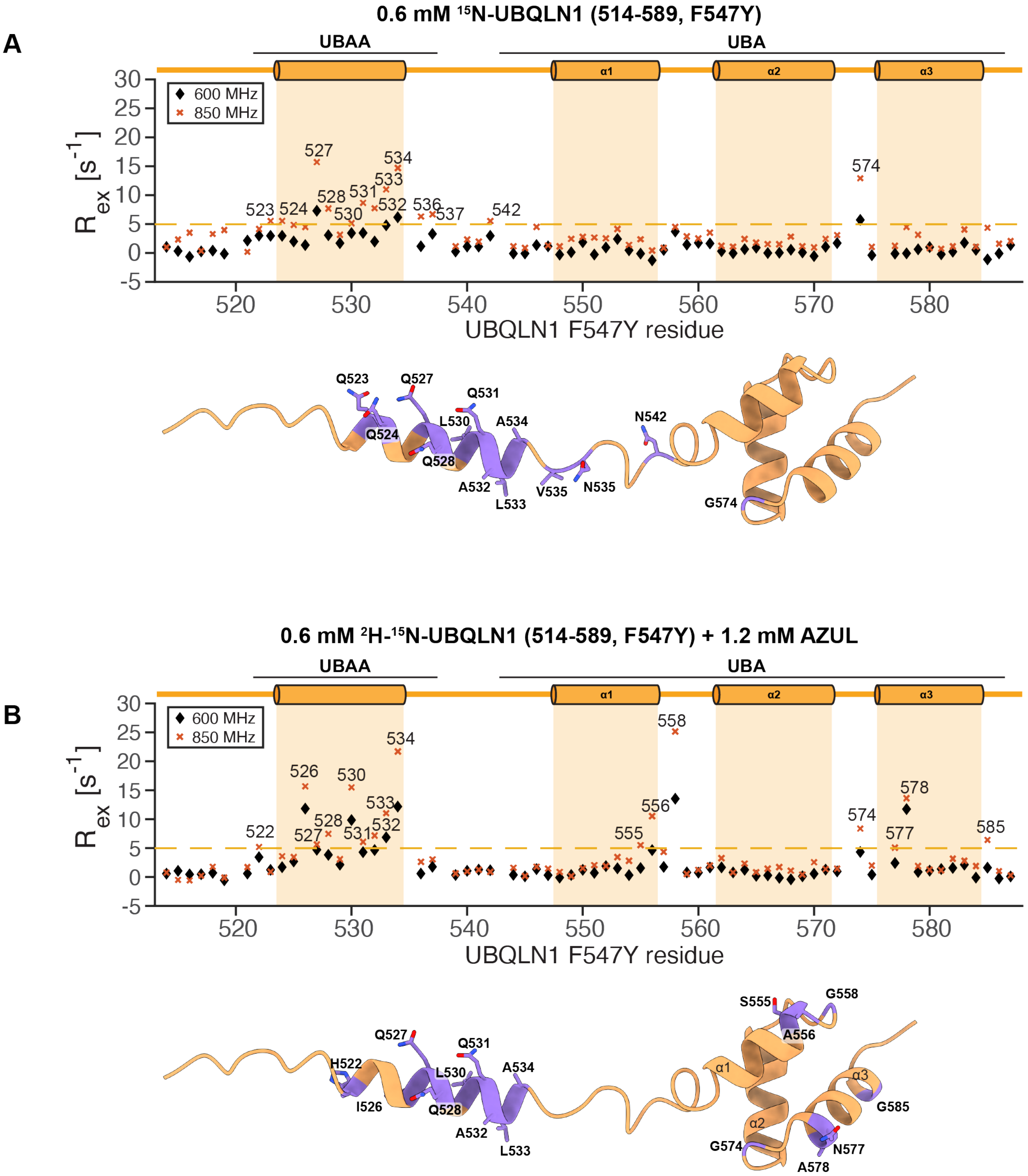


Figure S3. Related to Figure 4. A) $R_{ex}$ values from CMPG $R_2$ relaxation dispersion measurements acquired on 0.6 mM $^1$H-$^{15}$N-UBQLN1 (514-589, F547Y) at 850 or 600 MHz plotted according to UBQLN1 residue (top). $R_{ex}$ values above 5 $s^{-1}$ are shown in purple on the AlphaFold2-predicted structure of UBQLN1 514-589 (bottom). B) $R_{ex}$ values from CMPG $R_2$ relaxation dispersion measurements acquired on 0.6 mM $^2$H-$^{15}$N-UBQLN1 (514-589, F547Y) mixed with 1.2 mM AZUL at 850 or 600 MHz plotted according to UBQLN1 residue (top). $R_{ex}$ values above 5 $s^{-1}$ are shown in purple on the AlphaFold2-predicted structure of UBQLN1 514-589 (bottom).

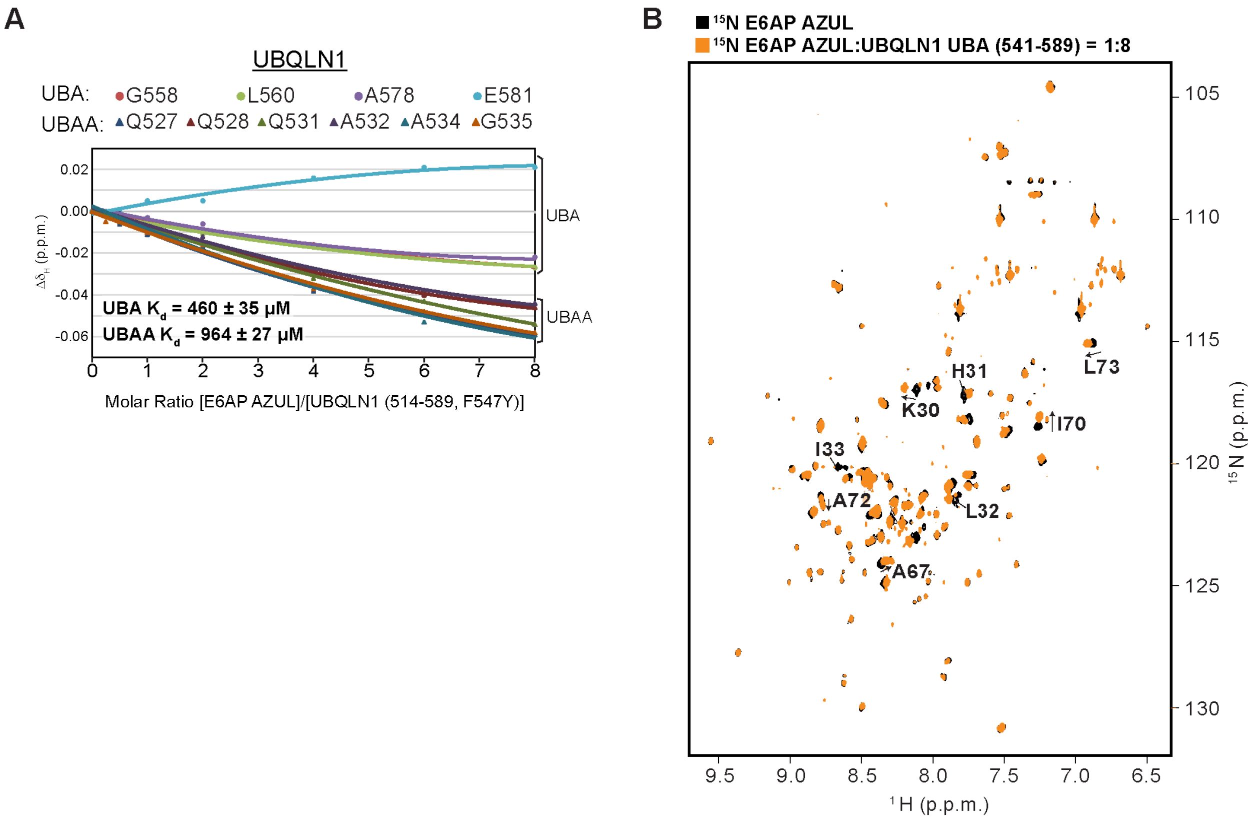


Figure S4. Related to Figure 5. A) Graph showing quantified hydrogen amide signal shifting of selected residues from the UBA or UBAA of 50 μM $^{15}$N-UBQLN1 (514-589, F547Y) at varying molar ratio with unlabeled AZUL, ending with a maximal concentration of 400 μM AZUL. The UBAA and UBA residues were separately fit to a one-site binding model as described in Methods to calculate a $K_d$ value based on their chemical shift changes. B) $^{15}$N-HSQC spectra of $^{15}$N-E6AP AZUL (black) overlayed with that of the $^{15}$N-E6AP AZUL mixed with 8-fold molar excess unlabeled UBQLN1 UBA 541-589 (orange). Signals for residues highlighted in Figure 2A are labeled.

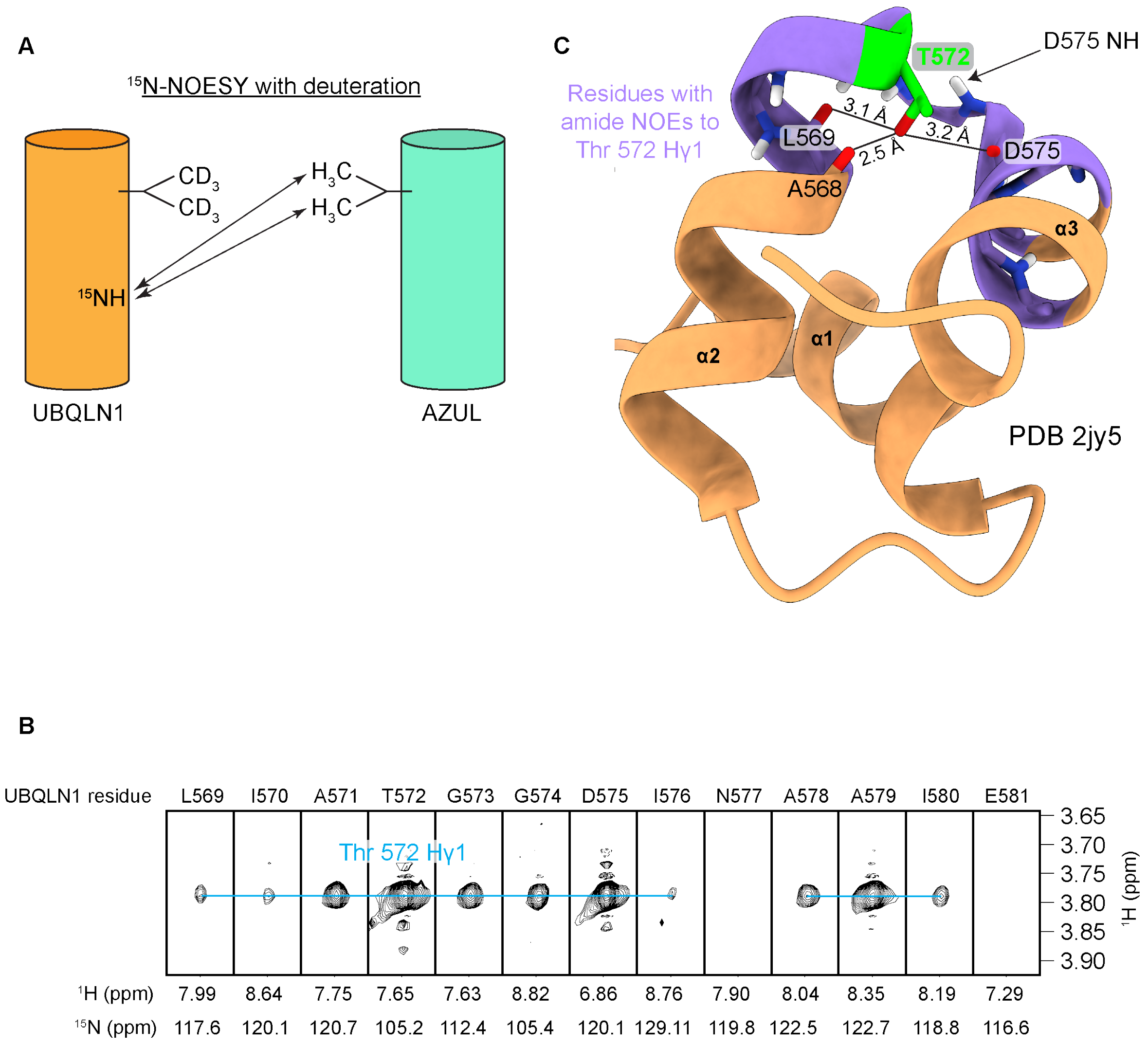
A
$^{15}$N-NOESY with deuteration
$CD_3$
$CD_3$
$H_3C$
$H_3C$
$^{15}$NH
UBQLN1
AZUL
C
Residues with amide NOEs to Thr 572 Hγ1
T572
D575 NH
3.1 Å
3.2 Å
2.5 Å
L569
D575
A568
α3
α2
α1
PDB 2jy5
B
UBQLN1 residue
L569 I570 A571 T572 G573 G574 D575 I576 N577 A578 A579 I580 E581
Thr 572 Hγ1
3.65
3.70
3.75
3.80
3.85
3.90
$^1$H (ppm)
$^1$H (ppm) 7.99 8.64 7.75 7.65 7.63 8.82 6.86 8.76 7.90 8.04 8.35 8.19 7.29
$^{15}$N (ppm) 117.6 120.1 120.7 105.2 112.4 105.4 120.1 129.11 119.8 122.5 122.7 118.8 116.6

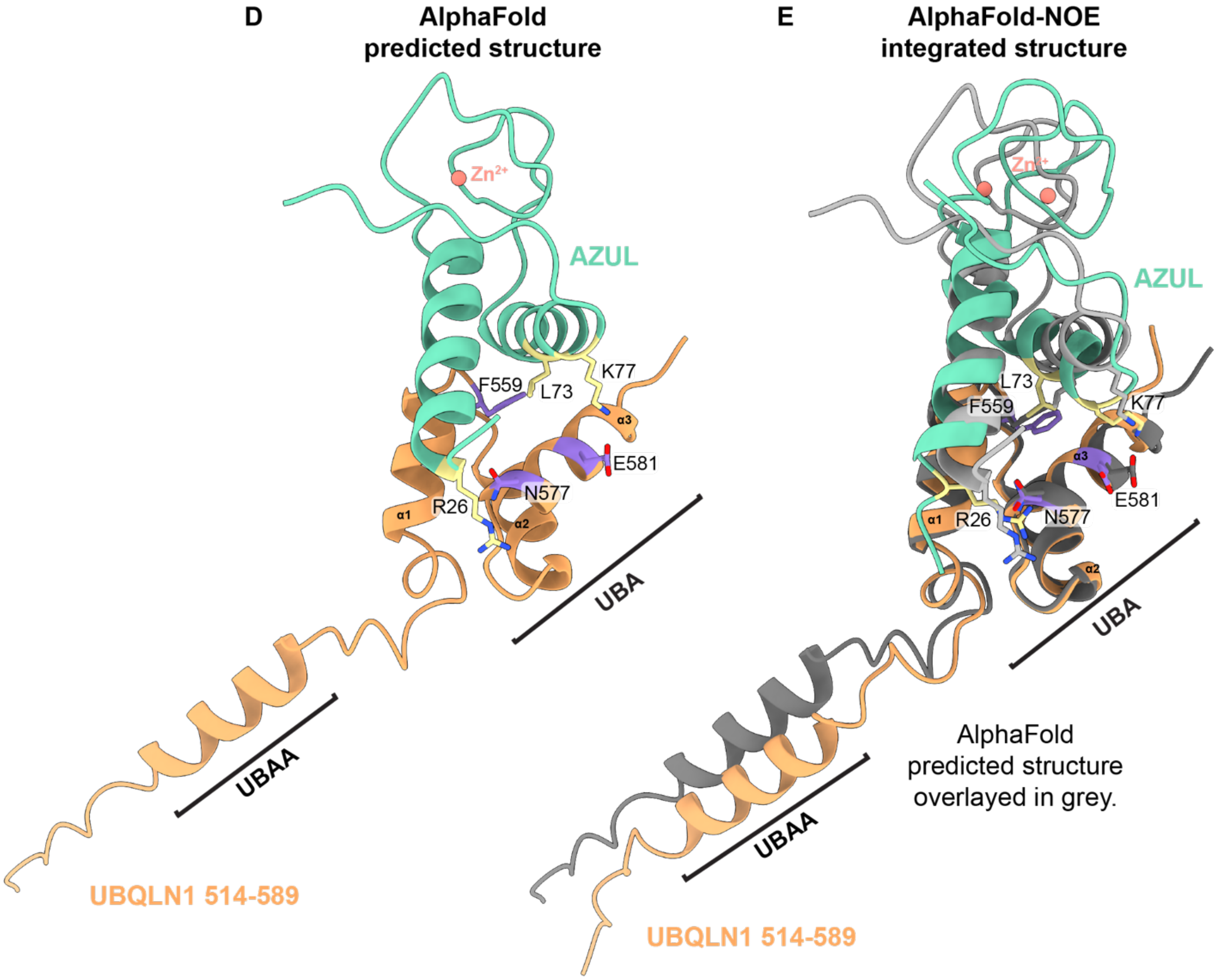


**F**

Candidate interactor

↓

co-IP in cells
2D NMR validation of direct interaction
interaction in droplets

↓

Intermolecular NOEs

↓

AlphaFold2-Multimer prediction

↓

AlphaFold-guided NOE
assignments

↓

AlphaFold-NOE integrated
structural model

Figure S5. Related to Figure 6. A) Schematic showing labeling scheme to unambiguously select for intermolecular NOE interactions between the amide protons of UBQLN1 and non-exchangeable protons of AZUL by using a standard $^{15}N$-NOESY. B) Selected regions of a $^{15}N$-edited NOESY collected on 0.6 mM $^{2}H$-$^{15}N$ UBQLN1 (514-589, F547Y) mixed with 1.2 mM unlabeled AZUL displaying NOEs from T572H$\gamma$1. C) Structure of the UBQLN1 UBA (PDB 2jy5) showing residues with NOEs from T572H$\gamma$1 in purple. Distance measurements from O$\gamma$1 to adjacent backbone oxygens are shown, indicating likely hydrogen bond acceptors. (D) AlphaFold2-Multimer-predicted structure of E6AP AZUL in complex with UBQLN1 (514-589). E) AlphaFold-NOE integrated structure overlayed with the AlphaFold2-Multimer-predicted structure (grey). In (A-B), residues from UBQLN1 with NOEs detected to their amides are shown in purple; residues from AZUL with assigned NOEs are shown in yellow. F) Workflow developed in the current study to use NMR and AlphaFold iteratively.
